# Independent evolution of atypical sperm morphology in a passerine bird, the red-browed finch (*Neochmia temporalis temporalis*)

**DOI:** 10.1101/2024.09.30.615754

**Authors:** Melissah Rowe, Daniel M. Hooper, Antje Hofgaard, Laura L. Hurley, Callum S. McDiarmid, Ioanna Pataraia, Jan T. Lifjeld, Simon C. Griffith

## Abstract

Spermatozoa exhibit striking morphological variation across the animal kingdom. In passerine birds, sperm exhibit considerable variation in size, yet the basic sperm phenotype is highly conserved; sperm are filiform, the head is corkscrew-shaped, and the midpiece is elongated and twisted around the flagellum. A significant departure from this typical sperm morphology has been reported in the sister species, the Eurasian bullfinch (*Pyrrhula pyrrhula*) and Azores bullfinch (*P. murina*). Here, we report a second evolutionary shift in passerine sperm phenotype in the nominate subspecies of the red-browed finch (*Neochmia temporalis temporalis*); sperm are non-filiform, with an ellipsoid head, and an extremely short midpiece restricted to the nuclear-axoneme junction. Additionally, we show that the sperm phenotype of the red-browed finch is similar to the putatively neotenous sperm described in the two bullfinch species. Using whole-genome data, we found no evidence that the unusual sperm phenotype of the red-browed finch is associated with reduced genetic variation or a population bottleneck. In contrast, we find some support for the hypothesis that relaxed post-copulatory sexual selection may, at least in part, explain the unusual sperm of the red-browed finch. We also discuss the possible roles of mutation, genetic drift, and genetic hitchhiking, in the evolutionary origins and maintenance of neotenous sperm phenotypes. Finally, we suggest that these dramatic evolutionary shifts in sperm phenotype warrant further investigation and highlight the need for a greater understanding of the developmental and genomic basis of sperm phenotype.

## Introduction

Sperm cells exhibit remarkable levels of variation in size, shape, and gross morphology across the animal kingdom. Indeed, sperm are the most morphologically diverse cell type known, with variation observed across all levels of organisation: among species, among populations of the same species, among males within a population, as well as both among and within ejaculates from a single male (Pitnick et al. 2009). This extraordinary diversity is hypothesized to be shaped by three main factors: (1) fertilization mode (i.e., internal vs. external fertilization; Franzén 1956; Kahrl et al. 2021, 2022), (2) post-copulatory sexual selection (i.e., sperm competition and cryptic female choice; Snook 2005; Simmons and Fitzpatrick 2012; Lupold and Pitnick 2018; Lüpold et al. 2020), and (3) phylogeny (Jamieson et al. 1995; Supriya et al. 2016; Omotoriogun et al. 2020). Yet, despite centuries of sperm biology research (Birkhead and Montgomerie 2009), understanding the rapid and divergent nature of sperm evolution remains an outstanding challenge in evolutionary biology.

Sperm morphological variation has been studied extensively in birds, with particular focus on the order Passeriformes; data from the passerine birds comprise more than 13% of all publicly available data on sperm morphology across the animal kingdom (https://spermtree.org; Fitzpatrick et al. 2022). In the songbirds (Passeriformes: Passeri), total sperm length varies considerably, ranging from approximately 43 μm to 290 μm (Pitnick et al. 2009; Omotoriogun et al. 2020). Despite variation in sperm length, songbird sperm have a conserved general structure; sperm are filiform, the head (i.e., acrosome and nucleus) is corkscrew-shaped, and the midpiece is elongated and wrapped helically around the flagellum (Jamieson 2007). In many songbird species, sperm also possess a helical membrane (also referred to as an acrosome keel) that spirals around the acrosome core (Jamieson 2007), though the extent of this helical keel is highly variable across species (Støstad et al. 2018). Of the more than 750 passerine species studied to date, only two species are noted as exhibiting atypical sperm morphology - the Eurasian bullfinch (*Pyrrhula pyrrhula*) and the Azores bullfinch (*P. murina*). In these species, mature sperm cells have an ellipsoid head with a rounded, not pointed, cap-like acrosome, and an extremely short midpiece (Birkhead et al. 2006; Lifjeld et al. 2013).

In both bullfinch species, ultrastructural studies also reveal several traits that make the sperm of these species distinct from sperm cells from other passerines. These include a simple rounded acrosome, poorly condensed chromatin in the nucleus, and a midpiece comprised of several small mitochondria (instead of a single fused, elongated, and helical mitochondrion) clustered around the nuclear-axonemal junction (Birkhead et al. 2007; Lifjeld et al. 2013). This sperm structure is suggested to resemble that of an immature sperm (i.e., spermatozoa neoteny) and to result from suppression of the final stages of spermiogenesis (Birkhead et al. 2007). Importantly, the identification of the Eurasian and Azores bullfinch as sister taxa (Töpfer et al. 2011), suggests a single ancestral origin of this unusual sperm morphology (Lifjeld et al. 2013).

Two main hypotheses have been proposed to explain the unusual sperm morphology of the bullfinches. First, Durrant et al. (2010) hypothesized that the highly variable and unusual sperm morphology of the Eurasian bullfinch may be the result of inbreeding or reduced genetic diversity following a bottleneck event. However, analysis of microsatellite data for this species provided no support for this hypothesis. Specifically, Durrant et al. (2010) compared levels of genetic diversity in the Eurasian bullfinch to three other fringillid finch species and found no differences in microsatellite polymorphism or heterozygosity among the species and no evidence for recent bottlenecks in any of the species, indicating that a reduction in genetic variation could not account for the unusual sperm morphology of the Eurasian bullfinch. Second, it has been hypothesized that the neotenous sperm of the bullfinches is the result of relaxed selection on sperm morphology due to low levels or an absence of sperm competition (Birkhead et al. 2006; Lifjeld et al. 2013). Furthermore, under a scenario of low or absent sperm competition, where any functional sperm should be able to fertilize ova, skipping the final stages of spermatogenesis is expected to reduce reproduction costs incurred by males. Thus, the unusual sperm morphology of bullfinches has been interpreted as an adaptation to reduce costs of sperm production (Birkhead and Immler 2006; Birkhead et al. 2007; Lifjeld et al. 2013).

Atypical sperm morphology and structure has also been noted in rodents. Among the murid rodents (Rodentia: Muridae), atypical sperm head morphology (i.e., a non-falciform shape) has been reported in several lineages (Breed 1983, 1993, 1995; Thitipramote et al. 2011). Most notably, the red veld rat (*Aethomys ineptus*) exhibits an unusual sperm head shape (rounded, not falciform), atypical acrosome structure, and a high degree of variability in chromatin condensation in the nucleus (Breed 1995, 1997). Similarly, in the naked mole rat (*Heterocephalus glaber,* Rodentia: Heterocephalidae), sperm are characterized by an atypical head shape and irregular and variable chromatin condensation in the nucleus (van der Horst et al. 2011). Importantly, such instances of extreme evolutionary shifts in sperm phenotype can provide novel insight into the evolutionary and developmental processes responsible for the diversification of sperm form and function.

Here, we report a second evolutionary shift in sperm phenotype in a passerine bird, the red-browed finch (*Neochmia temporalis*). Like the bullfinches, the red-browed finch exhibits a highly atypical sperm morphology. Our study has three main objectives. First, we wanted to know if the unusual sperm morphology of the red-browed finch is the same (or similar) to the unusual sperm phenotype of the Eurasian and Azores bullfinches. We therefore examined the sperm morphology (sperm size and general structure) of red-browed finches using brightfield microscopy and scanning electron microscopy (SEM) and examined sperm ultrastructure using transmission electron microscopy (TEM). Second, we compared the sperm morphology of the red-browed finch to that observed in several closely related Estrildidae finches using both published data (i.e., long-tailed finch, *Poephila acuticauda acuticauda*, Rowe et al. 2015b; masked finch, *P. personata*, McCarthy et al. 2021; zebra finch, *Taeniopygia castanotis*, Immler et al. 2012) and collection of sperm from two additional species (i.e., crimson finch, *N. phaeton*; double-barred finch, *Stizoptera bichenovii*). Notably, we also compared the sperm morphology of the nominate subspecies of red-browed finch (*N. temporalis temporalis*) to the lesser red-browed finch (*N. temporalis minor*). Among the passerines, the estrildid finches stand out as having particularly high levels of among-male variance in sperm morphology, likely the result of relaxed or weak selection on sperm morphology due to low levels of sperm competition in the clade (McCarthy et al. 2021). Thus, by comparing sperm morphology in several related taxa, we aim to gain insight into the evolution of the atypical sperm phenotype observed in the red-browed finch. Third, we investigated the hypothesis that the unusual sperm morphology observed in the red-browed finch is associated with reduced genetic variation or a population bottleneck. Specifically, using whole genome data, we directly compared the genetic diversity and demographic history of the red-browed finch with four estrildid finches, including taxa known to display ‘typical’ passerine sperm morphology.

## Materials and Methods

### Study species and sampling

The red-browed finch is a small (8.4-14.6 g), seed-eating Australian grassfinch (Estrildidae) endemic to the east coast of Australia (Payne 2020). Two subspecies are currently recognised: the nominate subspecies *N. temporalis temporali*s (distributed along the east coast, from Cooktown south to Victoria and South Australia and inland to western slopes of the Great Dividing Range and, in the south, west to Mt Lofty Ranges and Kangaroo Island); and *N. temporalis minor*, referred to as the lesser red-browed finch (distributed along the coast of northeast Queensland from Cape York Peninsula south to the Atherton Tablelands) (Payne 2020). A third subspecies has been intermittently recognized: *N. temporalis loftyi* (distributed in the southwest corner of South Australia around the Mt. Lofty Ranges) (Morcombe 2003); although this clade is currently placed within the nominate subspecies (Payne 2020). The red-browed finch is considered common throughout its range, with contemporary population size estimated to be in excess of 3 million individuals (Callaghan et al. 2021), and is a common aviary bird in Australia and globally.

We obtained sperm from a single male red-browed finch (*N. t. temporalis*) sampled at Lake Samsonvale (27°16′ S, 152°51′ E), 30km northwest of Brisbane (Queensland, Australia), during a field expedition in 2015. Subsequent light microscopy and SEM of the sperm from this individual revealed the unusual sperm morphology of the species. We therefore undertook targeted sampling across several locations in New South Wales and Queensland to obtain additional sperm samples from both wild and captive populations of red-browed finch. The aim of this sampling was to confirm the unusual sperm phenotype and obtain data on sperm size in this species. We obtained sperm samples from wild birds from several locations in New South Wales, including (i) Martinsville (n = 17), (ii) Nelson (n = 9), (iii) Narara (n = 9), (iv) Burrendong Arboretum (n = 2), (v) Wellington Caves (n = 2), and (vi) Ballimore (n = 1). We also obtained sperm samples from the lesser red-browed finch by sampling captive males (n = 5) held by an aviculturist in Queensland. Similarly, we sampled sperm from a male crimson finch held by an aviculturist in NSW. Finally, we sampled sperm from a single, wild male double-barred (*Stizoptera bichenovii bichenovii*) finch caught at Wianamatta Nature Reserve in NSW. All wild birds were caught with mist nets and released again at the site of capture after sampling. Full details of sampling locations are provided in the supplementary materials (Supplemental table S1).

In addition to sampling wild and captive finches in Australia, we obtained red-browed finches (males and females) belonging to the nominate subspecies from aviculturists in the Netherlands. Birds were transported to the Netherlands Institute of Ecology (NIOO-KNAW) and housed in small aviaries (50 x 180 x 40 cm, HxLxD) in mixed-sex groups of 2-4 individuals. Aviaries were fitted with several wooden perches and natural branches, wicker nests and nesting material (hay, coconut fibre, feathers), and food and water provided *ad libitum*. During sampling periods, birds were maintained under breeding conditions; 14L:10D photoperiod, breeding diet consisting of seed mixture, supplemented with a mix of sprouting seeds, egg food, live mealworms, and fresh millet sprays, as well as the occasional piece of fruit (apple, orange) and mix of green vegetables (e.g. peas, kale). We sampled sperm from captive males to investigate sperm size and sperm ultrastructure (see below). In addition, we present data on sperm swimming speed in these captive red-browed finches in the supplementary materials (S1).

### Sperm morphology and ultrastructure

We measured sperm size from the samples collected from red-browed finches across multiple populations in Australia (total n = 41), as well from sperm samples collected from captive Dutch birds (n = 3). We also measured sperm size from samples collected from captive male lesser red-browed finches (n = 5). Fresh sperm samples were collected via cloacal massage (Kucera and Heidinger 2018). Samples were immediately diluted in a small volume (∼ 10 μl) of phosphate buffered saline (PBS) before being fixed in 5% buffered formaldehyde solution. To quantify sperm size, we prepared sperm slides for brightfield microscopy. Specifically, a *c.* 20 μl aliquot of the fixed sperm sample was pipetted onto a glass slide, left to air dry overnight, and then rinsed gently with distilled water the next day before again being left to air dry. Next, we captured digital images of intact sperm cells and measured sperm length using SpermSizer v1.6.6 (McDiarmid et al. 2021). For the red-browed finch, we obtained measures for the length of the sperm head (including the acrosome, nucleus, and midpiece, as it was not possible to distinguish the midpiece using brightfield microscopy) and flagellum. For the lesser red-browed finch, we obtained measures for the length of the sperm head (acrosome and nucleus), midpiece length, and flagellum length. For both species, total sperm length was calculated as the sum of head and flagellum lengths. Following previous studies on sperm length (Laskemoen et al. 2007), we aimed to measure 10 sperm per sample. However, this was not always possible as many sperm samples contained very few sperm cells. In these instances, we quantified sperm length from as many cells as possible. For the red-browed finch, we measured an average of 8.77 ± 5.17 (mean ± SD) sperm cells (median: 8.0, range: 1-23). We tested for differences in sperm size (sperm head length, flagellum length, and total sperm length) among the different populations of the nominate subspecies of red-browed finch using ANOVA using R (v4.4.1; R Core Team 2024) and R studio (Posit team 2023). For the lesser red-browed finch, we measured an average of 12.60 ± 2.88 (mean ± SD) sperm cells (median: 12.0, range: 9-16).

We assessed intra-specific variation in sperm size for both the red-browed finch and the lesser red-browed finch by calculating the among-male and within-male coefficients of variation (Sokal and Rohlf 1981) in sperm length. Specifically, we calculated an adjusted among-male coefficient of variation in total sperm length as CVam = ((SD/X) x 100) x (1+(1/4N)), where SD is the standard deviation of mean total sperm length, X is mean total sperm length, and N is the number of males measured. Next, we calculated an adjusted within-male coefficient of variation (CVwm) of total sperm length using the same formula, though in this instance N is the number of cells measured per male.

Next, we examined sperm morphology (i.e., general structure) of the red-browed finch (n = 3), lesser red-browed finch (n = 4), crimson finch (n = 1), and double-barred finch (n = 1) using SEM. Briefly, 10 μl aliquots of formaldehyde-fixed sperm were pipetted onto glass coverslips precoated with poly-lysine (1mg/ml; Sigma P1274) and incubated overnight in a wet chamber at room temperature. Next, sperm adhering to the coverslips were dehydrated using a graded ethanol series (50, 70, 80, 90, 96 and 4 x 100% ethanol) and critical point dried (BAL-TEC CPD 030 Critical Point Dryer). Coverslips were then mounted on SEM stubs using carbon tape, sputter-coated with 4-6 nm platinum using a Cressington 308R coating system, and sperm cells imaged using a Hitachi S-4800 Field Emission Scanning Electron Microscope operated at 5.0 kV.

We examined sperm ultrastructure in the red-browed finch using TEM. To obtain enough sperm for TEM, we examined sperm within the seminal glomera using males from the captive Dutch population. Specifically, five males were euthanised (via anaesthetic overdose) and their seminal glomera isolated via dissection. Seminal glomera tissue was fixed in picric acid for 25 mins at room temp (4% paraformaldehyde, saturated picric acid, 1x PHEM buffer [pH 7.4]) and then stored in the fridge in 500 μl of 2% glutaraldehyde/4% paraformaldehyde/1x PHEM buffer [pH 7.4] until further processing. Samples were then washed 3x in 1x PHEM, postfixed with 1% OsO4 + ferricyanide followed by 3 washes in 1x PHEM. Samples were stained with 1% UA and finally washed another 3x with 1xPHEM. To infiltrate samples with epoxy, the samples were first dehydrated in a graded ethanol series at room temp (10 mins each in 50, 70, 80, 90, 96, and 4x 15 mins 100%) followed by infiltration with a EPON (without DMP30): ETOH 1:1, rotating over 2 days at room temperature. Subsequently, samples were further infiltrated with 100% EPON (with 1.5% DMP-30) and incubated overnight (rotating) at room temperature. Following infiltration, samples were placed in moulds and polymerized over two days at 60 °C. Finally, ultrathin sections (80 nm) were obtained with a Leica ultramicrotome equipped with a diatome diamond knife and observed using a JEOL 1400plus transmission electron microscope operated at 120 kV.

For the five males euthanised from the captive Dutch population, we also measured body mass (± 0.01 mg), dissected out the left and right testis, and immediately snap froze testis tissue before storing it at −80 °C. We later thawed the left testis and measured testis length and width (± 0.01 mm) using digital callipers. From these dimensions we calculated the mass of the left testis following (Møller 1991; Møller and Briskie 1995), estimated combined testes mass by doubling this value, and estimated relative testes mass for each male (testes mass/body mass x 100). We are aware that it would have been better to measure fresh testes mass or to obtain length/width data for both testes. However, this was not possible at the time of our sampling, and the approach used still provides an indication of relative testes size, and thus sperm competition, for the red-browed finch.

### Genomic sequencing, mapping, and genotyping

We generated whole genome sequence (WGS) data for the two subspecies of the red-browed finch (*N. t. temporalis*, N = 1; *N. t. minor*, N = 1) and two closely related Australian grassfinches belonging to the same subfamily (Poephilinae; family Estrildidae): the crimson finch (N = 1) and the double-barred finch (N = 2). Museum vouchered tissue samples were provided by the Australian National Wildlife Collection Genomic DNA was extracted using a DNeasy Blood & Tissue Kit (Qiagen, CA, USA) following the manufacturer’s protocol. WGS libraries were prepared for each sample using an Illumina TruSeq DNA Library Preparation Kit (Illumina, CA, USA) and sequenced on three lanes of an Illumina HiSeq2500 (2×150bp) and three lanes of an Illumina HiSeq 4000 (2×150 bp) at the University of Chicago Genomics Facility. We supplemented this genomic dataset with WGS data from previously published work for an additional four species in the family Estrildidae. WGS data for zebra finch (N = 3), long-tailed finch (N = 3), and double-barred finch (N = 1) were generated by Singhal et al. (2015) (ENA PRJEB10586). These three species belong to the same subfamily as the red-browed finch (Poephilinae; Olsson and Alström 2020). WGS data for the Bengalese finch (*Lonchura striata domestica*, N = 3) was generated by Lu et al. (2022) (NCBI PRJNA779480) and used for phylogenetic inference as an outgroup taxon within the Estrildidae (subfamily Lonchurinae).

Read data from each sample was trimmed of sequencing adapters and low-quality bases with cutadapt v4.8 (Martin 2015) before being mapped to the zebra finch reference genome v1.4 (GCF_003957565.2; Rhie et al. 2021) using BWA mem (Li 2013). PCR duplicates were removed using Picard’s MarkDuplicates module (http://broadinstitute.github.io/picard/). We recovered an average of 136.9 million paired end reads and a mean depth of coverage of 24.4× for the 15 samples included in this study (Supplemental table S2).

Genotyping was performed in accordance with GATK v4.4 Best Practices (van der Auwera and O’Connor 2020). In brief, an initial set of variants were called for each sample for each chromosome using HaplotypeCaller in GVCF move, GVCF output from all 15 samples were combined by chromosome using CombineGVCFs, and joint genotyping was performed using GenotypeGVCFs. We restricted downstream analyses to the 30 largest chromosomes in the zebra finch reference, together representing 98.3% of the genome, because we detected a roughly two-fold reduction in mapping performance and subsequent variant calling for the 10 smallest chromosomes <2.7 Mb in length. We selected all SNPs on each chromosome with the SelectVariants module of GATK, masked sites that overlapped with annotated repetitive regions of the zebra finch reference genome as called by repeatModeler2 (Flynn et al. 2020), and kept those that remained after applying the following variant quality filters: QD < 2, FS > 60.0, MQ < 30.0, ReadPosRankSum < −8.0.

### Phylogenetic analyses

We used vcftools v0.1.16 (Danecek et al. 2011) to generate a set of repeat-masked quality-filtered SNP variants from the 30 largest chromosomes separated by at least 10 kb, with a minor allele frequency of at least 2, and no missing genotype data across all samples. This pipeline produced 87,408 SNPs, 79,312 of which (those that had a minor allele in homozygosity in at least one individual) could be used for building a maximum likelihood tree with RAxML v8.2.4 (Stamatakis 2014). We implemented the ASC_GTRGAMMA model in combination with the Lewis correction for SNP ascertainment bias and carried out 350 bootstrap replicates. The three Bengalese finch samples were used as an outgroup in accordance with their established phylogenetic placement in previous analyses (Olsson and Alström 2020).

### Genetic diversity and demographic history

We evaluated evidence of reduced genetic variation or a population bottleneck in the red-browed finch by comparing both subspecies against the other four species in our genomic dataset belonging to the subfamily Poephilinae. We used bcftools v1.9 (Danecek et al. 2021) to count the number of heterozygous genotypes observed on each of the 30 largest chromosomes in each sample (Supplemental table S3). We calculated individual heterozygosity as the number of heterozygous sites divided by the number of nucleotide positions in that sample with a minimum depth of coverage of 10 for each chromosome. Heterozygosity was then evaluated as a single summary statistic by summing values across the 29 largest autosomal chromosomes. Under the assumption that the unusual sperm morphology of the red-browed finch is a result of reduced genetic diversity, we expect genomic heterozygosity to be lowest in the red-browed finch, with larger heterozygosity values observed in all other finch taxa.

We next applied the Pairwise Sequential Markovian Coalescent (PSMC; (Li and Durbin 2011) to each genome to model how effective population sizes (N_e_) have changed over evolutionary time. For this analysis, we only considered the 29 largest autosomal chromosomes >2.7 Mb in length. Diploid consensus sequences were generated for each sample using a lower depth of coverage threshold of 10 and an upper depth of coverage threshold equal to twice the mean depth of coverage for that sample. Default parameters were used for PSMC module fq2psmcfa, except that we used a quality filter of 20 and a bin size of 50 bp, to account for the higher density of heterozygous sites in finches compared to humans. Adopting the recommendations of Hilgers et al. (2024), we adjusted default tuning for the number of time intervals to avoid false population size peaks/crashes and ran PSMC with the following parameters: -N30, -t5, -r5, -p “2+2+25*2+4+6”. We converted evolutionary units to millions of years and N_e_ using a generation time of 2 years (Bird et al. 2020) and a germline mutation rate of 5.85e^-09^ (per site, per generation) based on pedigree inferred germline mutation rate for the zebra finch (Bergeron et al. 2023). We note that the inferred absolute values of N_e_ and number of generations are highly dependent on our assumptions about the mutation rate and generation time. We therefore focus our evaluation on the relative differences between taxa as these qualitative trends (e.g. a population bottleneck) are not affected by parameter tuning.

## Results

All male red-browed finches from the nominate subspecies, including both wild and captive birds, exhibited sperm morphology that is highly atypical for passerines; sperm were small, with an ellipsoid, ‘tadpole-like’ head and lacked an elongated midpiece. Across all males, total sperm length was 39.14 ± 3.88 μm (mean ± SD), with a size range of 30.73-46.85 μm, which is considerably shorter than sperm size reported in other estrildid species (table 1). Total sperm length was highly variable both within- and among-males (table 1). Notably, the coefficient of variation in total sperm length among-males was 9.96%, which is on par with values reported for the bullfinches (Lifjeld et al. 2013) and among the highest reported for passerine birds (McCarthy et al. 2021). Total sperm length did not differ among the populations (F_7,36_ = 1.72, p = 0.14). Similarly, neither sperm head length nor sperm flagellum length differed among the populations (head: F_7,36_ = 0.91, p = 0.51; flagellum: F_7,36_ = 1.59, p = 0.17). Considering all males sampled (n=5), relative testes mass was 0.18 ± 0.14% for captive male red-browed finches. However, in 1 of the 5 males dissected, we were unable to obtain a sperm sample at the time of testes sampling, suggesting that this male might not have been reproductively active. When we excluded this male, relative testes mass was 0.21 ± 0.13%.

**Table 1.** Sperm size in the red-browed finch and related Australian estrildid finches, including the lesser red-browed finch (this study), long-tailed finch (data from Rowe et al. 2015b), masked finch (data from McCarthy et al. 2021), and zebra finch (data from Immler et al. 2012); Data for zebra finch taken from NSW wild population, midpiece length is reported as straight midpiece length, which is a calculated measure and longer than standard midpiece length reported for the other species, CVam calculated from available data, ^‡^ reported in (McCarthy et al. 2021). Data shown are mean ± SD.

| Species | Head length<br>($\mu\text{m}$) | Midpiece<br>length<br>($\mu\text{m}$) | Flagellum<br>length<br>($\mu\text{m}$) | Total sperm<br>length ($\mu\text{m}$) | CVwm | CVam |
| --- | --- | --- | --- | --- | --- | --- |
| Red-browed finch (n = 44) | 7.39 ± 1.4 | NA | 31.75 ± 3.7 | 39.14 ± 3.9 | 14.77 | 9.96 |
| Lesser red-browed finch (n=5) | 12.71 ± 1.2 | 5.08 ± 0.7 | 45.05 ± 4.2 | 57.75 ± 5.3 | 7.33 | 9.54 |
| Long-tailed finch | 12.80 ± 0.7 | 39.98 ± 4.4 | 62.70 ± 4.6 | 75.50 ± 4.4 | 2.71 | 5.82 |
| Masked finch | 10.83 ± 1.0 | 25.21 ± 4.7 | 49.61 ± 3.3 | 60.34 ± 3.5 | 5.75 | 5.9 |
| Zebra finch | 11.9 ± 0.6 | 32.2 ± 4.2 | 54.2 ± 4.8 | 66.0 ± 4.8 | 6.7-7.3<sup>†</sup> | 7.3 |

In contrast, under brightfield microscopy, sperm of the lesser red-browed finch appeared to have typical passerine sperm morphology, albeit with an extremely short midpiece (table 1). In the lesser red-browed finch, total sperm length was 57.75 ± 5.3 μm (mean ± SD), with a size range of 54.16-66.86 μm. Thus, the lesser red-browed finch has longer sperm than the nominate subspecies of the red-browed finch, but, on average, shorter sperm compared to the other estrildid finches examined (table 1). Total sperm length was also highly variable within- and among-males in the lesser red-browed finch (table 1).

Scanning electron microscopy (SEM) confirmed that the sperm morphology of the nominate subspecies of red-browed finch differs markedly from the typical sperm morphology observed in most passerines. Specifically, sperm of the red-browed finch are non-filiform, with an ellipsoid head incorporating a rounded, cap-like acrosome and lacking a helical membrane, and an extremely short midpiece restricted to the base of the head (Figure 1A, 1B). In contrast, sperm of the lesser red-browed finch possess the distinctive corkscrew-shaped head typical of passerine sperm (Figure 1C), and both the crimson finch and the double-barred finch exhibit highly typical passerine sperm morphology; a corkscrew-shaped head with an acrosomal keel and an elongated midpiece wrapped helically around the flagellum (Figure 1D, 1E). In the red-browed finch (nominate subspecies), we also observed considerable variation in the overall shape of the sperm head, both among- and within-males (Figure 2). Intriguingly, in the lesser red-browed finch, SEM revealed an apparent lack of an acrosomal keel on the corkscrew-shaped sperm head and confirmed the presence of a very short midpiece wrapping helically around the flagellum (Figure 1C, Figure S1).

**Figure 1.**
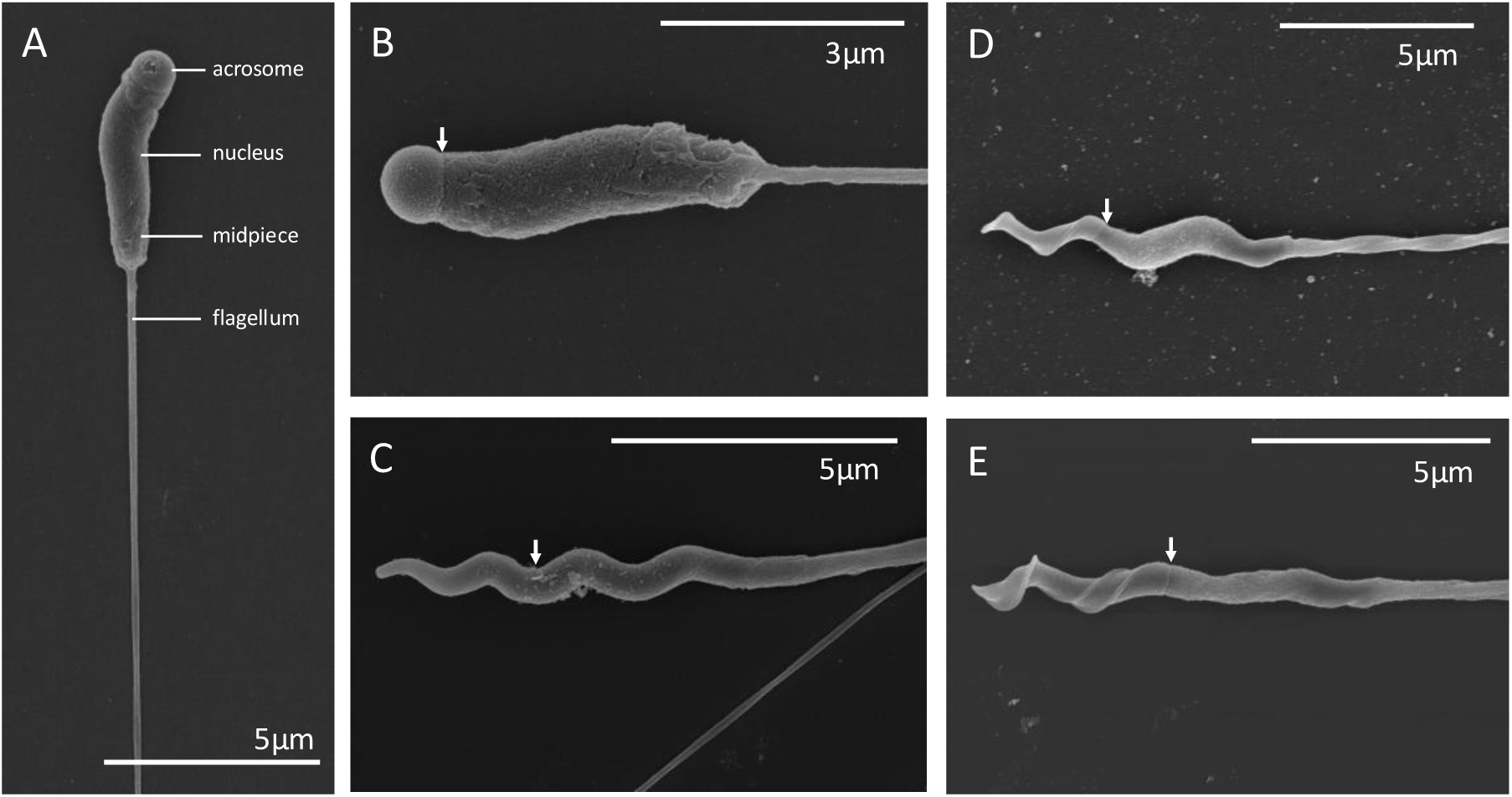
Scanning Electron micrographs of sperm cells from Australian finches. (A) Sperm from the red-browed finch showing the sperm head (acrosome and nucleus), the midpiece, and the anterior portion of the flagellum (which clearly lacks an elongated midpiece). Sperm head morphology of (B) the red-browed finch, (C) lesser red-browed finch, (D) crimson finch, and (E) double-barred finch. Arrows indicate the junction between the acrosome and nucleus.

**Figure 2.**
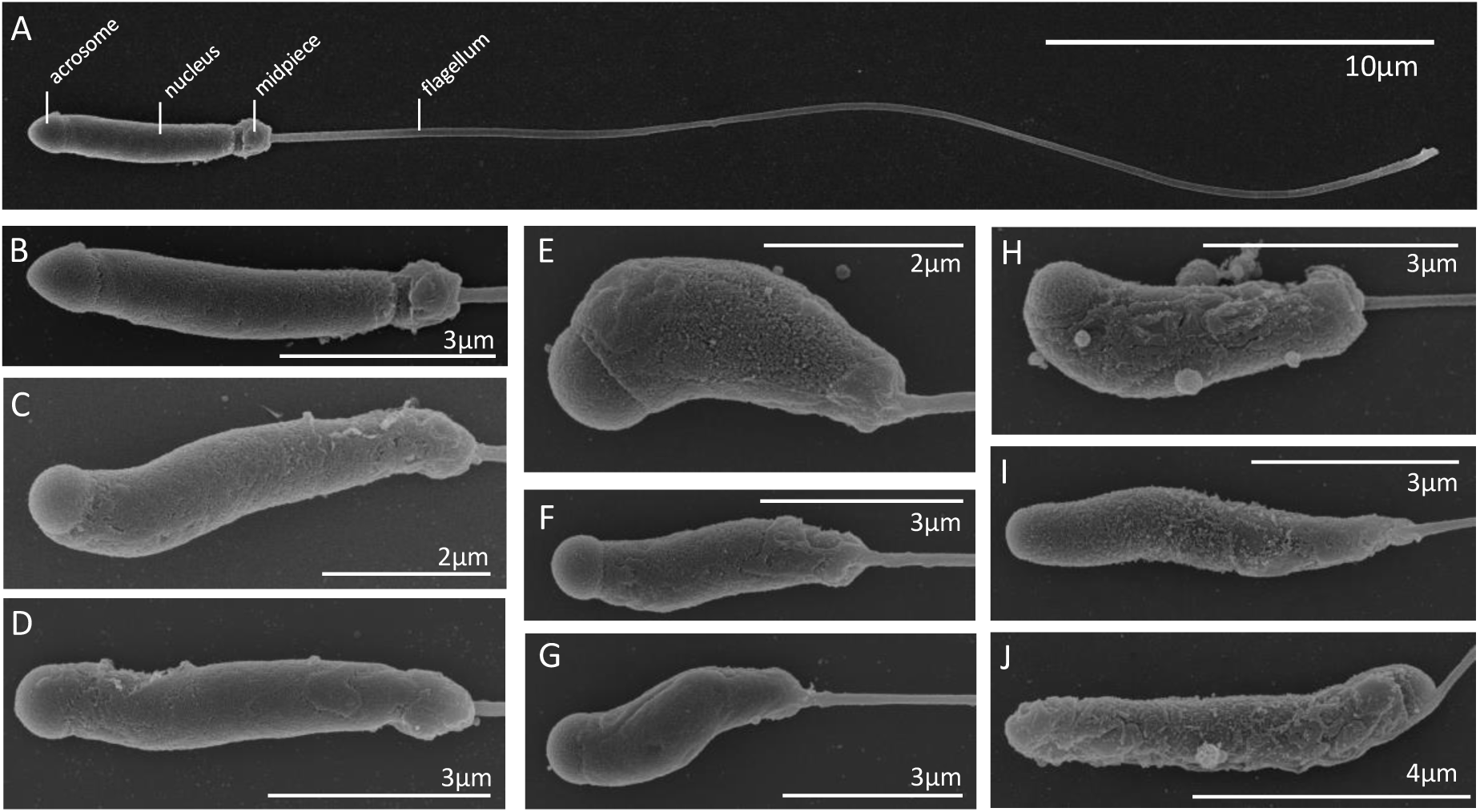
Scanning electron micrographs of the atypical sperm of the red-browed finch. (A) Overall structure of a sperm cell showing the sperm head (acrosome and nucleus), midpiece, and flagellum. Note the midpiece is clustered at the base of the nucleus and does not extend along the flagellum. (B-J) Variation in sperm head morphology within-males (i.e. sperm from the same ejaculate) and among-males (A-D male 1; E-G male 2; H-J male 3).

Transmission electron microscopy of sperm from the nominate subspecies of red-browed finch confirmed the presence of multiple mitochondria profiles clustered at the base of the nucleus (nuclear-axonemal junction) and a rounded sperm head with a cap-like acrosome (Figure 3). Sperm exhibited considerable variation in the density of chromatin within the nucleus, with some cells showing regions of strong chromatin condensation (Figure 3A, 3B), while other cells showed more heterogeneity in chromatin density throughout the nucleus, including the occurrence of chromatin-less regions (Figure 3C). Finally, TEM images suggest there may be a reduction in the size of the outer dense fibres of the sperm axoneme of the red-browed finch (Figure S2).

**Figure 3.**
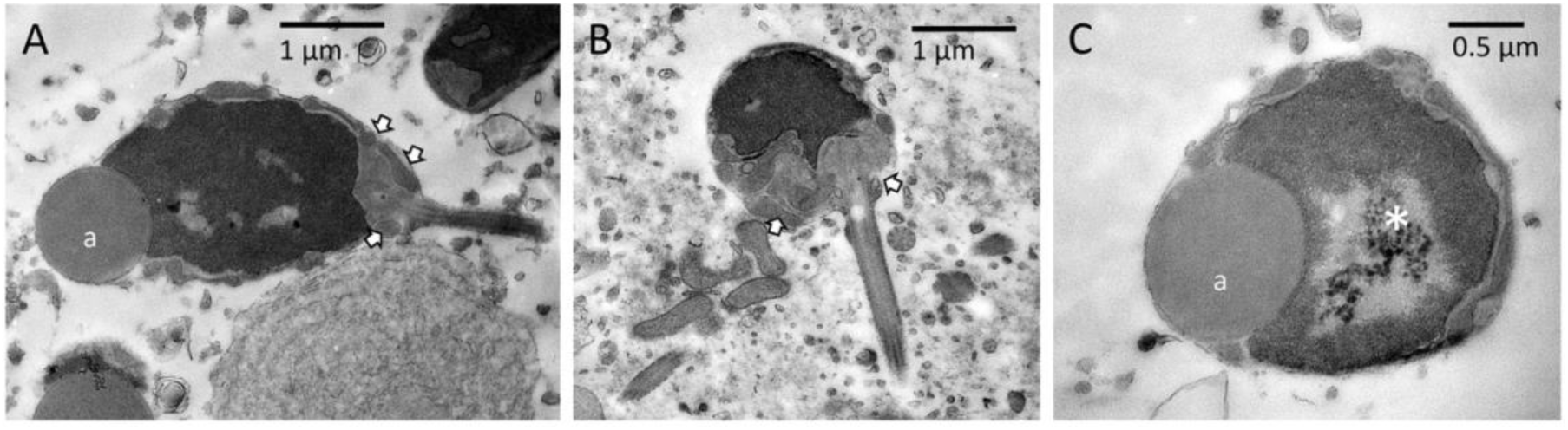
Transmission electron micrographs of sperm cells from the red-browed finch. (A) sperm head with a rounded shape and a cap-like acrosome (a), (B) sperm head with multiple mitochondria profiles clustered at the nuclear-axonemal junction, (C) sperm head with poor chromatin condensation and chromatin-less region (*). Arrows indicate mitochondria profiles clustered near the base of the sperm head.

Based on gross morphology and ultrastructure, the nominate subspecies of red-browed finch appears to have the same atypical sperm phenotype as the Eurasian and Azores bullfinch. In these three species, sperm possess a rounded head and the midpiece is not elongated, but rather clustered at the base of the head; these features stand in stark contrast to sperm morphology in passerine birds, including closely related taxa such as the lesser red-browed finch and the European greenfinch (Figure 4, Figure S3).

**Figure 4.**
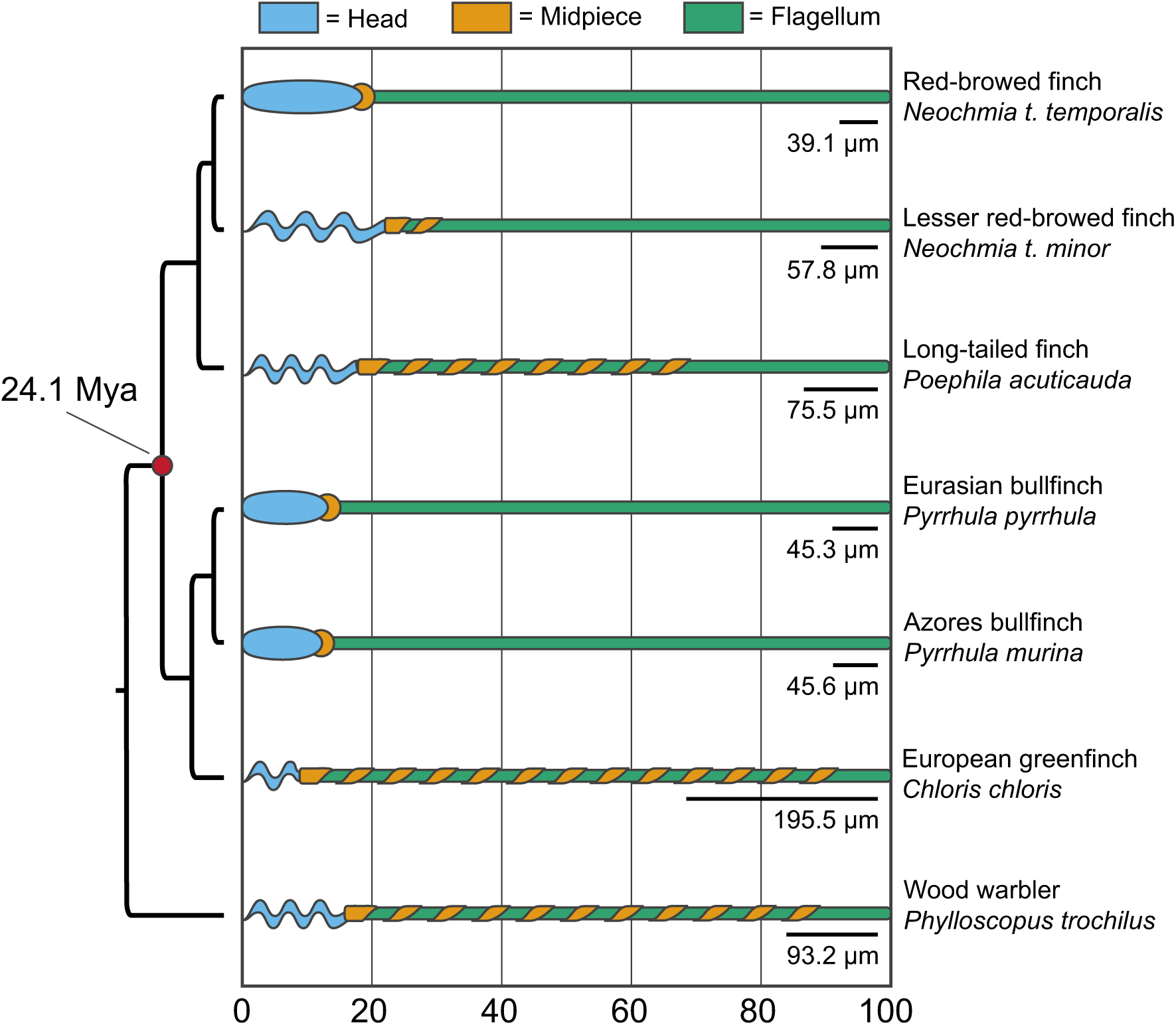
Stylized representations of sperm morphology in passerines showing the atypical sperm of the nominate subspecies of red-browed finch (*N. t. temporalis*), the Eurasian bullfinch (*P. pyrrhula*), and the Azores bullfinch (*P. murina*) compared to the typical sperm morphology of passerines, including closely related taxa within the Estrildidae (lesser red-browed finch [*N. t. minor*] and long-tailed finch [*P. acuticauda*]) and the Fringillidae (European greenfinch [*Chloris chloris*]), as well as a representative outgroup (Wood warbler [*Phylloscopus trochilus*]). Sperm drawings show the average percent of each sperm component for the species represented; for species with atypical sperm morphology there are no available measures of midpiece length, rather sperm head length represents the length of the head and midpiece combined. Black bars show total sperm length for each species. Phylogeny highlights the deep divergence between the Estrilidadae and Fringillidae and thus the independent origin of the unusual sperm phenotype. Sperm size data from this study, Lifjeld et al. (2013), and Rowe et al. (2015a,b).

Genetic diversity, as measured by mean site-based observed heterozygosity, varied considerably across finch taxa. The nominate subspecies of the red-browed finch displayed relatively low levels of heterozygosity compared to some of the other Australian grassfinch species evaluated. However, heterozygosity was even lower in both the lesser red-browed finch and the crimson finch (Figure 5, Supplemental table S4). The phylogenetic placement of finch taxa in the maximum-likelihood tree generated for this study is congruent with the relationships presented in the recent molecular phylogeny of (Olsson and Alström 2020). Specifically, the phylogenetic tree supports the sister clade grouping of the red-browed finch and the crimson finch (Figure 6A). We estimated historical effective population sizes (N_e_) using the PSMC approach for all estrildid finches examined over a range from 2 million years ago (mya) up until about 10 thousand years ago (kya). This allowed us to qualitatively compare population trends among finch taxa. We recovered considerable variability in maximum N_e_ among species (Figure 6), broadly consistent with the differences between them we observed in heterozygosity. Within species, we also observed considerable variation in N_e_ over time, suggesting that most populations have gone through periods of population expansion and contraction likely linked with Pleistocene glacial cycle, extinction of megafauna, and altered fire regimes (Rule et al. 2012). For example, the nominate subspecies of the red-browed finch had a period of population expansion from approximately 500 kya until 50 kya after which N_e_ has experienced a gradual decline; although historically the taxon has exhibited a similar N_e_ trajectory to the other finches examined (Figure 6B). Notably, the zebra finch has had a substantial and lasting period of demographic expansion, resulting in relatively large contemporary population sizes (Figure 6C). In contrast, the lesser red-browed finch has experienced a consistently low effective population size with signatures of a gradual and continued decline over the last 1 million years (Figure 6B). In the crimson finch (Figure 6F), a species displaying typical passerine sperm morphology, both historical and contemporary N_e_ are substantially lower than that observed in the nominate red-browed finch.

**Figure 5.**
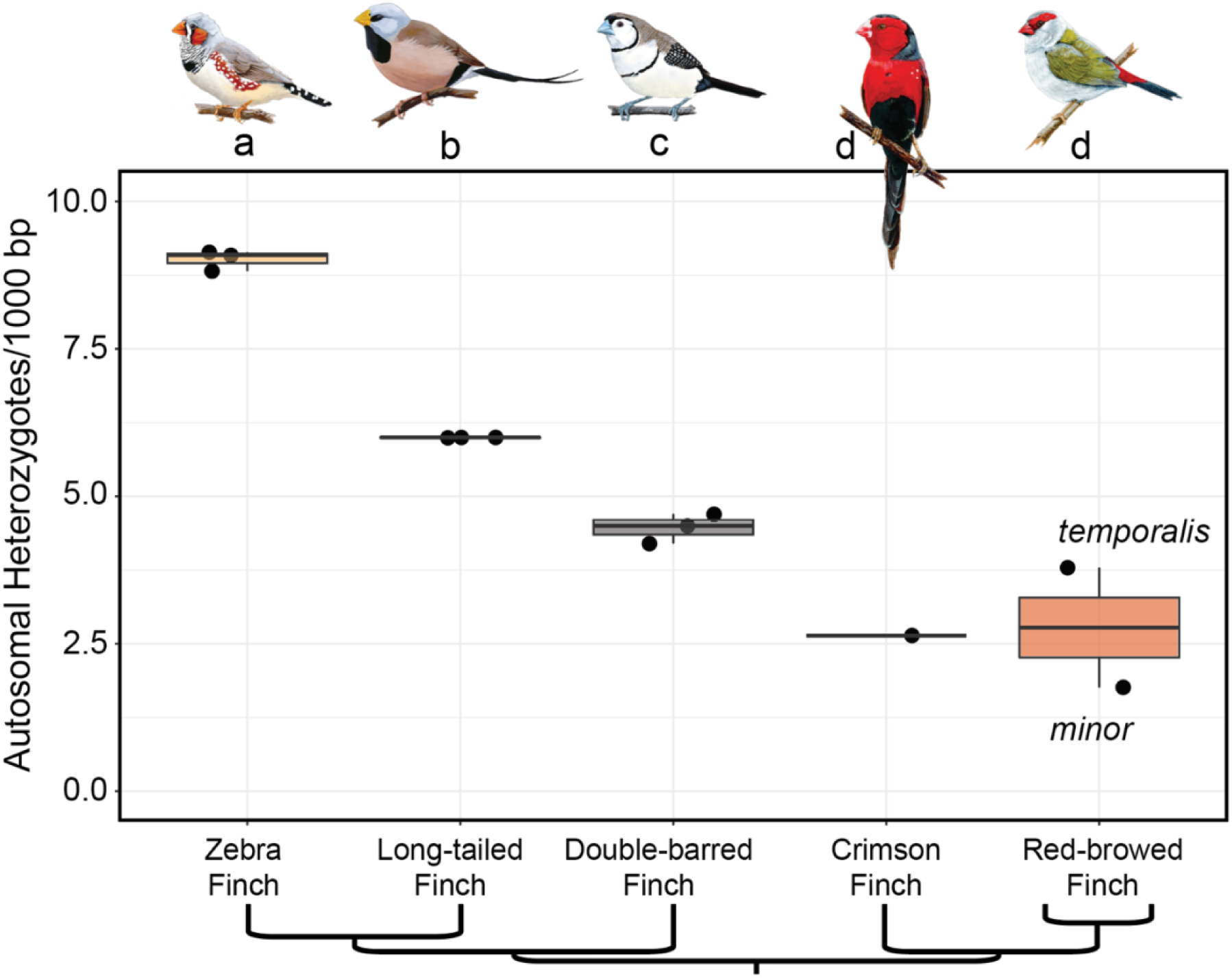
Mean site-based autosomal heterozygosity of six Australian grassfinch taxa, including the zebra finch, long-tailed finch, double-barred finch, crimson finch, and two subspecies of red-browed finch (the nominate subspecies, *N. t. temporalis* and the lesser red-browed finch, *N. t. minor*). Boxplots represent the variability within taxa and individuals are depicted as black points. Different letters beneath artistic renderings of each species indicate statistical significance between groups (adjusted P < 0.05, Tukey’s HSD test). A phylogeny beneath the panel depicts the relatedness among taxa.

**Figure 6.**
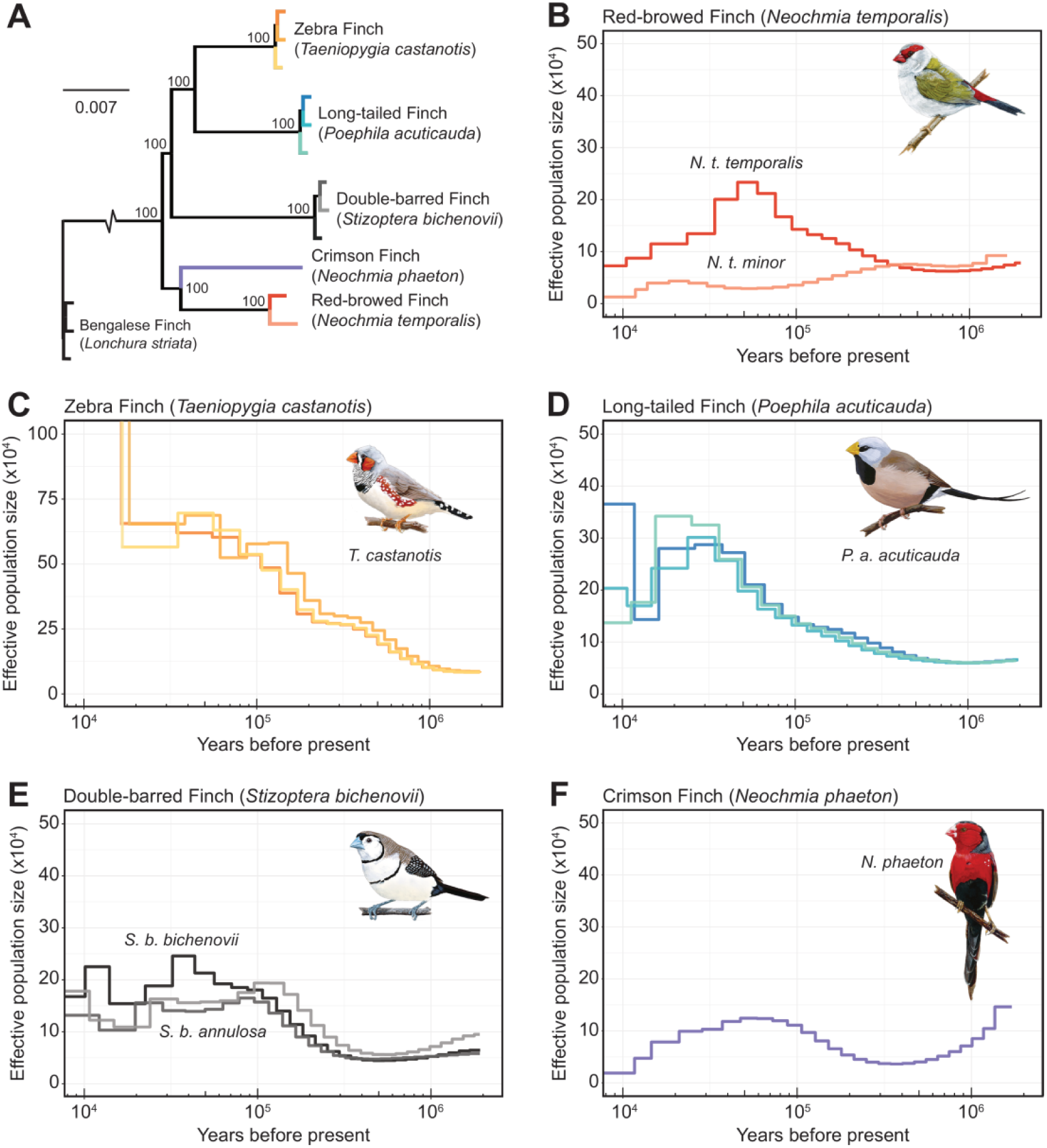
Phylogenetic and demographic history of the estrildid finches. (A) Maximum likelihood phylogenetic tree constructed with RAxML for five Australian grassfinch species. Support values of each node are based on 350 bootstrap replicates. The colors used for terminal branches in the phylogeny match an individual in the PSMC panels that follow. Inferences of demographic history for (B) two subspecies of red-browed finch (the nominate subspecies, *N. t. temporalis*, and the lesser red-browed finch, *N. t. minor*), (C) zebra finch, (D) long-tailed finch, (E) double-barred finch (subspecies *S. b. bichenovii* and *S. b. annulosa*), and the (F) crimson finch. We set the generation time (*g*) to 2 years and the germline mutation rate (μ) to 5.85e^-09^ per site per generation. Note that the y-axis scale bar differs between panel C and all other PSMC panels and that an inferred increase in N_e_ for the zebra finch in the last 20 thousand years has been omitted for clarity.

## Discussion

Sperm morphology is highly variable across species (Pitnick et al. 2009) and is frequently used as an informative trait for phylogenetic inference and taxonomy (Jamieson 1987; Jamieson et al. 1995; Bartolomaeus et al. 2024). In passerine birds, sperm are filiform and have a pronounced cork-screw shaped head and an elongated midpiece wrapped helically around the flagellum (Jamieson 2007). In stark contrast to this typical sperm phenotype, our study showed that the nominate subspecies of the red-browed finch exhibits highly atypical sperm morphology; sperm are non-filiform, with an ellipsoid head and an extremely short midpiece, consisting of multiple mitochondrial profiles, restricted to the nuclear-axonemal junction. Moreover, we found that the sperm of the red-browed finch appear very similar to the sperm morphology of the Eurasian bullfinch and Azores bullfinch (Birkhead et al. 2006, 2007; Lifjeld et al. 2013). Importantly, given that the split between the Estrildidae and the Fringillidae (bullfinches) dates to 24.14 mya (21.2 – 27.3, 95% HPD; Hooper and Price 2017), the unusual sperm morphology of the red-browed finch represents an independent evolutionary shift in sperm phenotype and only the second report of such a phenotypic shift in birds.

The sperm morphology of the nominate subspecies of the red-browed finch also stands out in comparison to the sperm of other Australian estrildid finches. In the crimson finch, the sister species to the red-browed finch, sperm exhibit typical passerine sperm morphology. Similarly, typical passerine sperm morphology is apparent in the double-barred finch (this study), long-tailed finch (Rowe et al. 2015b), masked finch (McCarthy et al. 2021), and zebra finch (Birkhead et al. 2005; Immler et al. 2012). Intriguingly, in the lesser red-browed finch, sperm possess the corkscrew-shaped head morphology typical of passerines. However, SEM revealed an apparent lack of an acrosomal keel in the lesser red-browed finch. Moreover, the midpiece, while elongated, is very short relative to the sperm of other passerines, including other estrildid finches. Thus, the lesser red-browed finch can be viewed as having a somewhat intermediate sperm morphology, though more comprehensive sampling and TEM would allow for a clearer understanding of sperm phenotype in this species. Notably, sperm morphology in the lesser red-browed finch appears similar to sperm collected from Beavan’s bullfinch (*P. erythaca*), a close relative of the Eurasian bullfinch (Birkhead et al. 2006). The two subspecies of red-browed finch are inferred to have diverged around 1.2 mya (Olsson and Alström 2020), and the shared reduction in midpiece length suggests that some degree of evolutionary change to sperm ontogeny may have pre-dated this split. In contrast, the highly atypical head shape of the nominate subspecies of red-browed finch appears to be a derived trait and represents a dramatic evolutionary shift in sperm phenotype.

Durrant et al. (2010) tested and rejected the hypothesis that reduced genetic diversity resulting from inbreeding following a bottleneck event might explain the unusual sperm morphology of the Eurasian bullfinch. Similarly, here, we find no evidence that reduced genetic diversity underlies the unusual sperm morphology of the red-browed finch; the inferred population demography of the red-browed finch does not indicate that the taxon has undergone a bottleneck event, and the subspecies exhibits a comparable level of heterozygosity to some of the other estrildid species we examined. The rationale underlying the reduced genetic diversity hypothesis is largely centred around findings that sperm abnormalities are more frequent in small, inbred populations or populations exhibiting recent reductions in heterozygosity (Gage et al. 2006; Roldan and Gomendio 2009). However, as highlighted by Lifjeld et al. (2013), the unusual sperm morphology of bullfinches are not sperm abnormalities, but instead reflect the typical sperm phenotype for these species.

Durrant et al. (2010) also speculated that reduced genetic diversity might explain the high levels of within- and among-male variation in sperm morphology observed in the Eurasian bullfinch. High levels of among-male variation in sperm length appears to be a feature of the estrildid finches (McCarthy et al. 2021). In the zebra finch, the remarkable among-male variation in sperm morphology is explained by a large polymorphic inversion on the Z chromosome (Kim et al. 2017; Knief et al. 2017). Estrildid finches appear particularly prone to inversions; karyotype structure is highly variable and appears to evolve rapidly, at least under certain demographic and geographic conditions (Christidis 1987; Hooper and Price 2015, 2017). Thus, an intriguing possibility is that chromosomal rearrangements – and especially those on the Z chromosome – might often underlie the high levels of sperm length variation in estrildid finches, though studies investigating the genetic architecture of sperm traits in a range of finch species are needed.

An alternate explanation for the unusual sperm morphology observed in the red-browed finch is that selection on sperm morphology is relaxed. Indeed, relaxed post-copulatory sexual selection has been suggested to explain neotenous sperm phenotypes observed in both birds and mammals (Birkhead and Immler 2006; Birkhead et al. 2006, 2007; Breed et al. 2007; Durrant et al. 2010; van der Horst et al. 2011; Lifjeld et al. 2013). Furthermore, given the sperm neoteny is thought to result from the skipping of the final stages of spermiogenesis, it has been posited that neotenous sperm may represent an adaptation to reduce the costs associated with sperm production when sperm competition is weak or absent (Birkhead and Immler 2006; Birkhead et al. 2007; Lifjeld et al. 2013). The red-browed finch appears to have extremely small testes, with relative testis mass estimated to be lower than those reported for the Eurasian bullfinch (0.22% and 0.29%, Birkhead et al. 2006; Lifjeld et al. 2013) and at the lower end of the range reported for passerine birds generally (0.14-9.75%, testes data from Møller 1991). This suggests that sperm competition is likely to be low in the red-browed finch. More generally, estrildid finches are thought to experience low levels of sperm competition (McCarthy et al. 2021) and available data on extra-pair paternity rates in this clade support this prediction (Brouwer and Griffith 2019). Thus, it seems plausible that relaxed selection on sperm phenotype may also help explain the neotenous sperm phenotype of the red-browed finch, although we suggest that consideration of additional hypotheses would be useful.

Understanding the evolutionary forces underlying the origin and maintenance of the neotenous sperm phenotype is challenging, not least because the phenomenon is so rare. Novel traits may spread to fixation within a population via genetic drift. In the nominate subspecies of red-browed finch there is no evidence of a population bottleneck and N_e_ has never been particularly low. Furthermore, the contemporary population size of the red-browed finch (all subspecies combined) is estimated at 3,594,643 individuals (Callaghan et al. 2021). Thus, it seems unlikely that genetic drift plays a role in the atypical sperm morphology in the red-browed finch. Alternatively, fixation of the unusual sperm phenotype may be the result of genetic hitchhiking; allele(s) underlying the neotenous sperm phenotype may go to fixation simply because they are in linkage disequilibrium with a second locus that is under selection (Maynard-Smith and Haigh 1974). Importantly, hitchhiking can explain the fixation of neutral alleles or in some cases mildly deleterious alleles (Maynard-Smith and Haigh 1974; Good and Desai 2014). Unfortunately, we currently have insufficient knowledge of the genetic basis of the unusual sperm morphology in red-browed finches, or indeed any taxa, and we lack the population level sampling necessary for understanding of the structure of linkage disequilibrium in this species. As such, we refrain from ruling out this hypothesis and suggest it warrants further consideration. Finally, assuming relaxed sexual selection (due to an absence of sperm competition), if skipping the final stages reduces the cost of sperm production and thus provides a fitness benefit to males, the trait may spread through adaptive evolution. Such a scenario is currently the leading hypothesis explaining the neotenous sperm phenotype across taxa, and is supported by evidence suggesting that sperm competition is low or absent in all taxa exhibiting this sperm morphology (Birkhead et al. 2006; Breed et al. 2007; van der Horst et al. 2011; Lifjeld et al. 2013), including the red-browed finch. It is worth noting, however, that several passerine species reported as having extremely low levels of extra-pair paternity (i.e. < 5% offspring classified as EPP) or to be genetically monogamous (i.e., 0% EPP, see Brouwer and Griffith 2019) display typical passerine sperm morphology, including the common redstart (*Phoenicurus phoenicurus*), Eurasian jackdaw (*Corvus monedula*), marsh warbler (*Acrocephalus palustris*), northern wheatear (*Oenanthe oenanthe*), white-throated dipper (*Cinclus cinclus*), common crossbill (*Loxia curvirostra*), canary (*Serinus canaria*), and wild zebra finch (Durrant et al. 2010; Immler et al. 2012; Støstad et al. 2018; NHMO Sperm collection, Lifjeld 2019). This suggests that a novel mutation, rather than standing genetic variation, may be critical to the emergence of the unusual and novel sperm phenotype.

There is some evidence in passerine birds that the purifying selection on genes acting during the final stages of spermatogenesis is relatively relaxed (Segami et al. 2022), suggesting that a loss-of-function mutation arising in genes involved in late-stage spermatogenesis may underlie the origins of the unusual sperm phenotype. A key feature of the atypical sperm phenotype is the rounded or ellipsoid shape of the head and the variability in the extent of chromatin condensation in the nucleus. In passerines, sperm chromatin condensation occurs during the last phase of spermatogenesis (i.e., spermiogenesis), with progressive compaction of chromatin associated with elongation of the sperm head (Aire 2014). Protamines are small, sperm-specific nuclear proteins that replace histones during the final stages of spermiogenesis and are crucial to the packaging and condensing of DNA within the sperm nucleus. Unfortunately our understanding of sperm protamines in passerine birds is limited (but see Nakano et al. 1976; Chiva et al. 1987; Oliva and Dixon 1989; Ausió et al. 1999 for studies of protamines in non-passerine birds). In mammals, however, alterations in protamine expression can have major effects on sperm phenotype and male fertility (Carrell and Liu 2001; Aoki et al. 2005; Lüke et al. 2014; Arévalo et al. 2022). For example, in mice, alterations to protamine expression results in unusual sperm head morphologies and poor chromatin condensation, the latter noted as being similar in appearance to the incompletely packed chromatin of spermatids (Cho et al. 2003). Thus, we hypothesize that changes in protamine genes or regulatory regions, or alternatively in accessory or transition proteins involved in chromatin condensation, may play a role in the unusual sperm head morphology reported previously in both mammals (Breed 1995, 1997; van der Horst et al. 2011) and bullfinches (Birkhead et al. 2006, 2007; Lifjeld et al. 2013), as well as here in the nominate subspecies of the red-browed finch.

In conclusion, our results demonstrate that the nominate subspecies of the red-browed finch displays highly atypical sperm for a passerine bird and that it shares this atypical phenotype with the Eurasian and Azores bullfinch. Although the evolutionary factors responsible for the unusual sperm phenotype remain to be determined, our findings suggest that relaxed or weak selection on sperm morphology may explain, at least in part, the unusual sperm morphology in the red-browed finch. However, we suggest that consideration of additional explanations, such as genetic hitchhiking, might help explain how the trait has spread to fixation, especially given the relatively large population size of the red-browed finch. More generally, we also suggest that studies investigating the genes associated with the unusual sperm phenotype, and how these genes influence sperm development during spermatogenesis, will provide much needed insight into the function and evolution of the neotenous sperm phenotype as well as contribute to a broader understanding of sperm morphological diversity across the animal kingdom.

## Data availability

Data on sperm size are archived on Dryad (doi:10.5061/dryad.73n5tb362). All genomic sequence data generated for this project will be archived through NCBI (BioProject ID: PRJNA1158757).

## Acknowledgements

We are most grateful to Mike Webster, Diana Carneiro, Bruce Hockley, Geoff Dennis, for assistance with fieldwork and sampling of wild finches, and to Shane Hauschildt and Mike Fidler for providing access to their captive estrildids. We are also grateful to Anne Dijkzeul, Nina Teeuw, and Ruben de Wit for caring for the finches held at the Netherlands Institute of Ecology, and to Nanneke van der Wal for assistance with animal ethics permission in The Netherlands. We thank Judith Risse and Becky Cramer for useful discussions, Allison Johnson for providing drawings of the different finch species included in this manuscript, and Lars Erik Johannessen and Gaute Grønstøl for assistance with sample management at the Natural History Museum Oslo (Norway). This study was supported by a Norwegian Research Council grant to MR (230434), an Australian Research Council Grant to SCG, DMH and MR (DP180101783), and a Norwegian Research Council grant to JTL (301592). Whole genome sequence data was generated using funding from a National Science Foundation DDIG award to DMH (1601323).

## Animal Ethics

All trapping and sampling of finches in Australia was conducted in accordance with local regulations and under license (in NSW: ARA 2017/032 and ARA 2020/020; in QLD: WISP15212314 and A2100). All import of samples from Australia to the Netherlands was done under permit (PWS2015-AU-002216 and PWS2021-AU-000562). All work undertaken with captive red-browed finches in the Netherlands was performed under appropriate license (AVD 80100 202115477) and approved by the Animal Experimentation Committee of the Royal Dutch Academy of Sciences (DEC-KNAW; protocol number NIOO23.06).

## Author Contributions

Conceptualization: M.R.; Data curation: M.R. and D.M.H.; Formal analysis: M.R. and D.M.H.; Investigation: M.R., D.M.H., A.H., and I.P.; Project administration: M.R.; Resources: M.R., D.M.H., A.H., L.L.H., C.S.M., J.T.L., and S.C.G.; Visualization: M.R., A.H., and D.M.H.; Writing – original draft: M.R. and D.M.H., Writing – review & editing: M.R., D.M.H., A.H., L.L.H., C.S.M., I.P., J.T.L., and S.C.G.

## Supplementary Materials

### S1. Sperm swimming speed in the red-browed finch

#### Materials and methods

We quantified sperm swimming speed using fresh sperm samples collected from six captive red-browed finch males (Netherlands captive birds). Sperm samples were obtained by cloacal massage (Kucera and Heidinger 2018) and immediately diluted in ∼50 μl PBS. After mixing gently, 6 μl of the diluted sample was loaded onto a chamber slide (depth 20 μm, Leija, Netherlands) placed on a heating stage and sperm swimming speed assessed by computer-assisted-sperm-analysis (CASA) using the Sperm Class Analyzer ® (SCA, Microptic, Barcelona). All buffers and equipment were preheated to 40°C (avian body temperature). Sperm recordings were captured at 50 frames per second using negative phase contrast microscopy at 100x magnification with a Basler acA1300-200uc camera connected to a Nikon E200 microscope. We recorded up to 9 fields of view (range: 6-9) to maximise the number of sperm cells tracked; all recordings were completed within 1-minute of sample loading. All recordings were later visually examined, and any tracking or detection errors (e.g. detection of cell debris) deleted manually. We considered sperm immotile when with VCL < 20.0 μm/s (based on analysis of dead sperm videoed using the same capture settings to identify the kinematic profile of drifting sperm). For each male, we recorded an average of 78.8 ± 93.5 (mean±SD) motile sperm cells (median: 41.5, range: 5– 242); although half of the males had a low number of motile sperm cells (i.e., < 20) in their samples, we report mean sperm swimming speed across all males sampled following recommendations of (Cramer et al. 2019). From these analyses, we obtained data on curvilinear velocity (VCL), straight line velocity (VSL), average path velocity (VAP) of the sperm cells.

#### Results

The average sperm swimming speed (*VCL*) from captive male red-browed finches was 93.26 ± 8.84 μm/s (mean±SD; *VAP*: 82.96 ± 14.44; *VSL*: 74.85 ± 13.66), which is within the range previously reported for passerines species (VCL 77-164 μm/s; (Kleven et al. 2009) and sits between measures reported for the Eurasian bullfinch (i.e., 21.65 μm/s (Birkhead et al. 2006); 157.3 μm/s (Lifjeld et al. 2013).

## Supplementary Figures

**Figure S1.**
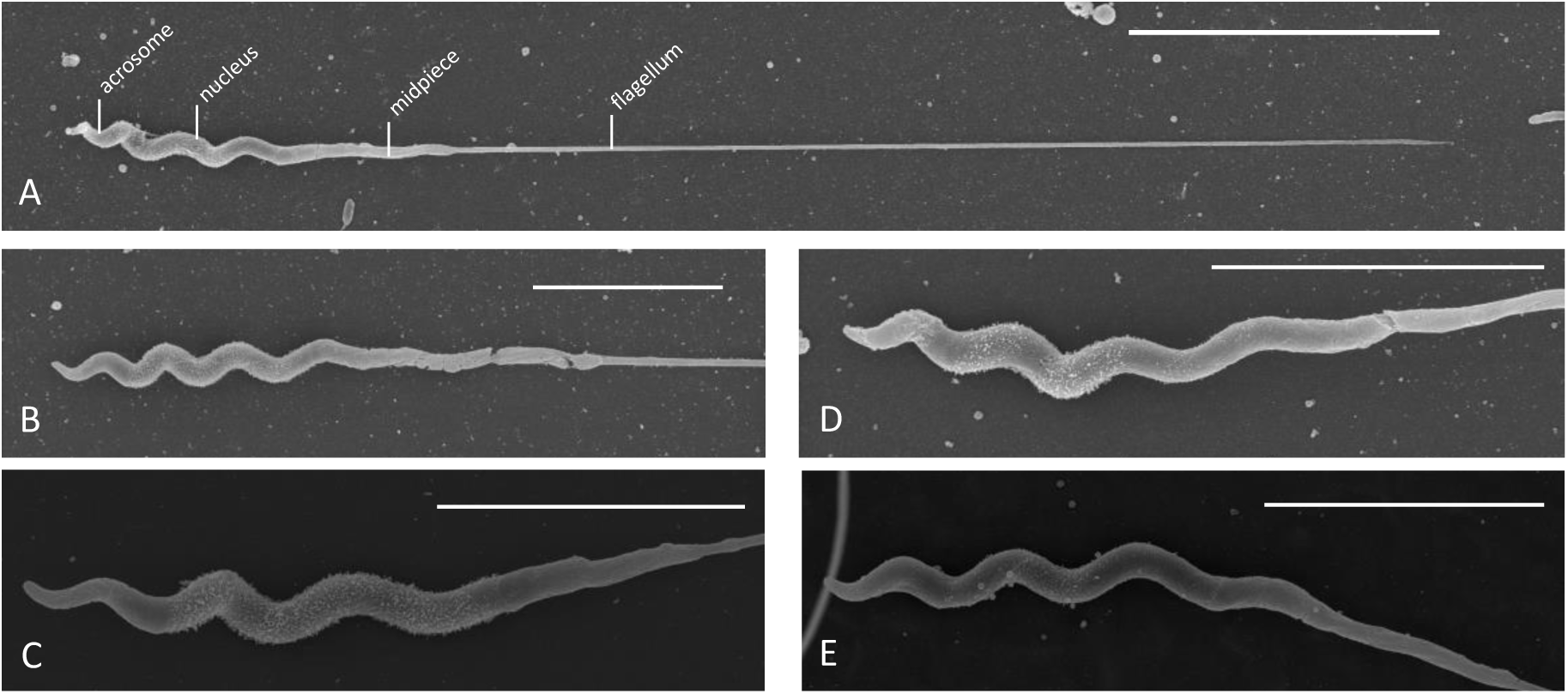
Scanning electron micrographs of the sperm of the lesser red-browed finch (*N. t. minor*). (A) Overall structure of a sperm cell showing the sperm head (acrosome and nucleus), midpiece, and flagellum. (B-E) Variation in sperm head morphology. Note the midpiece is elongated, but relatively short in length. Scale bar = 10 μm in A; 5 μm in B-E.

**Figure S2.**
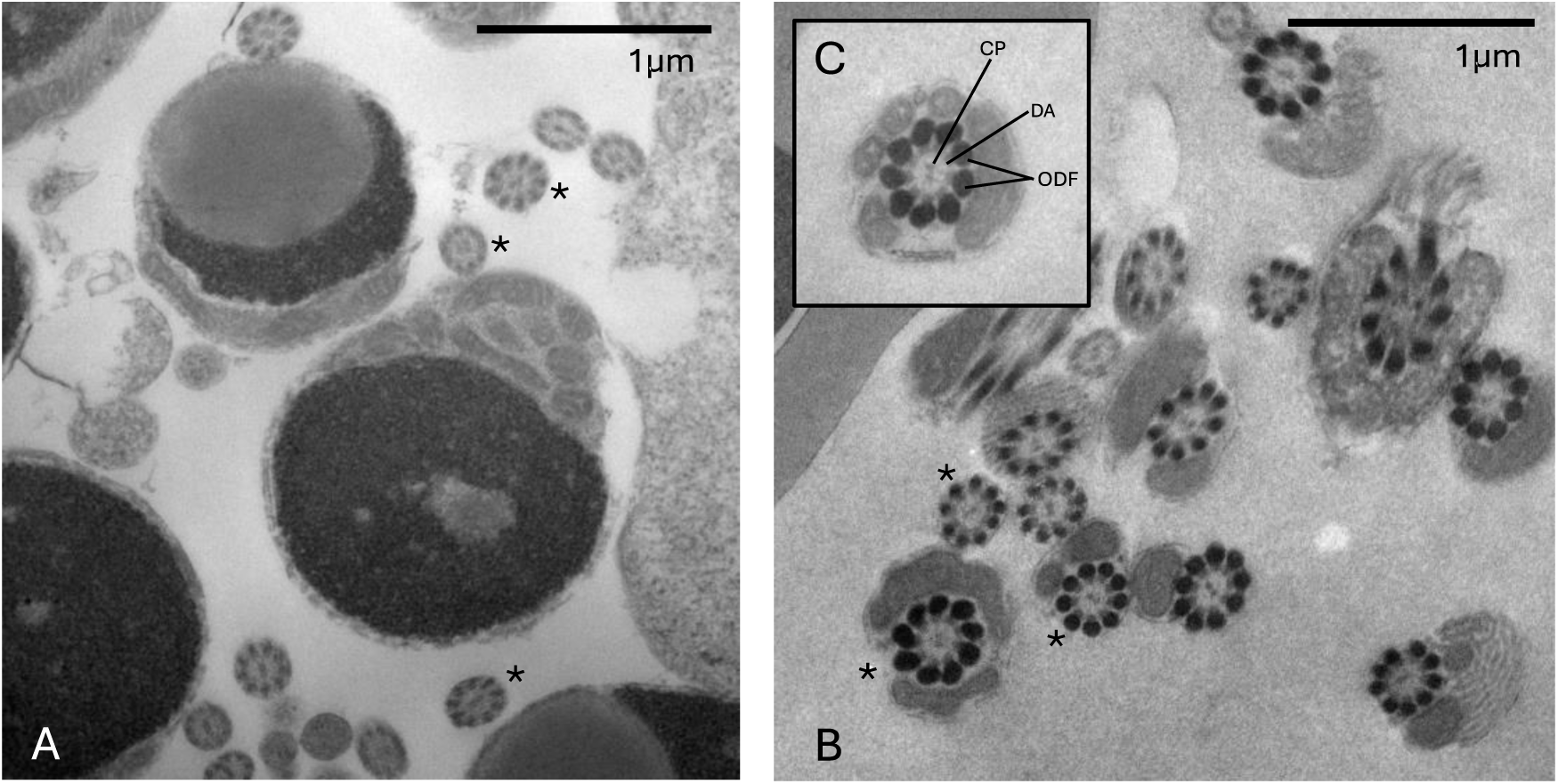
Transmission electron micrographs showing transverse sections of the flagella axoneme of the (A) red-browed finch (*N. t. temporalis*) and (B) long-tailed finch (*P. a acuticauda*, Rowe unpublished). (C) detail of the flagella axoneme showing the central microtubule pair (CP), the inner dynein arms (DA), and outer dense fibres (ODF). The microtubule pair and dynein arms form the archetypical 9 + 2 microtubule structure of the axoneme (two central microtubule singlets and nine outer microtubule doublets). * identify flagella sections in both species showing variation in the size of outer dense fibres (ODF), note that the variation and overall size of the ODF appear greater in long-tailed finch (shown in B) relative to the red-browed finch (shown in A).

**Figure S3.**
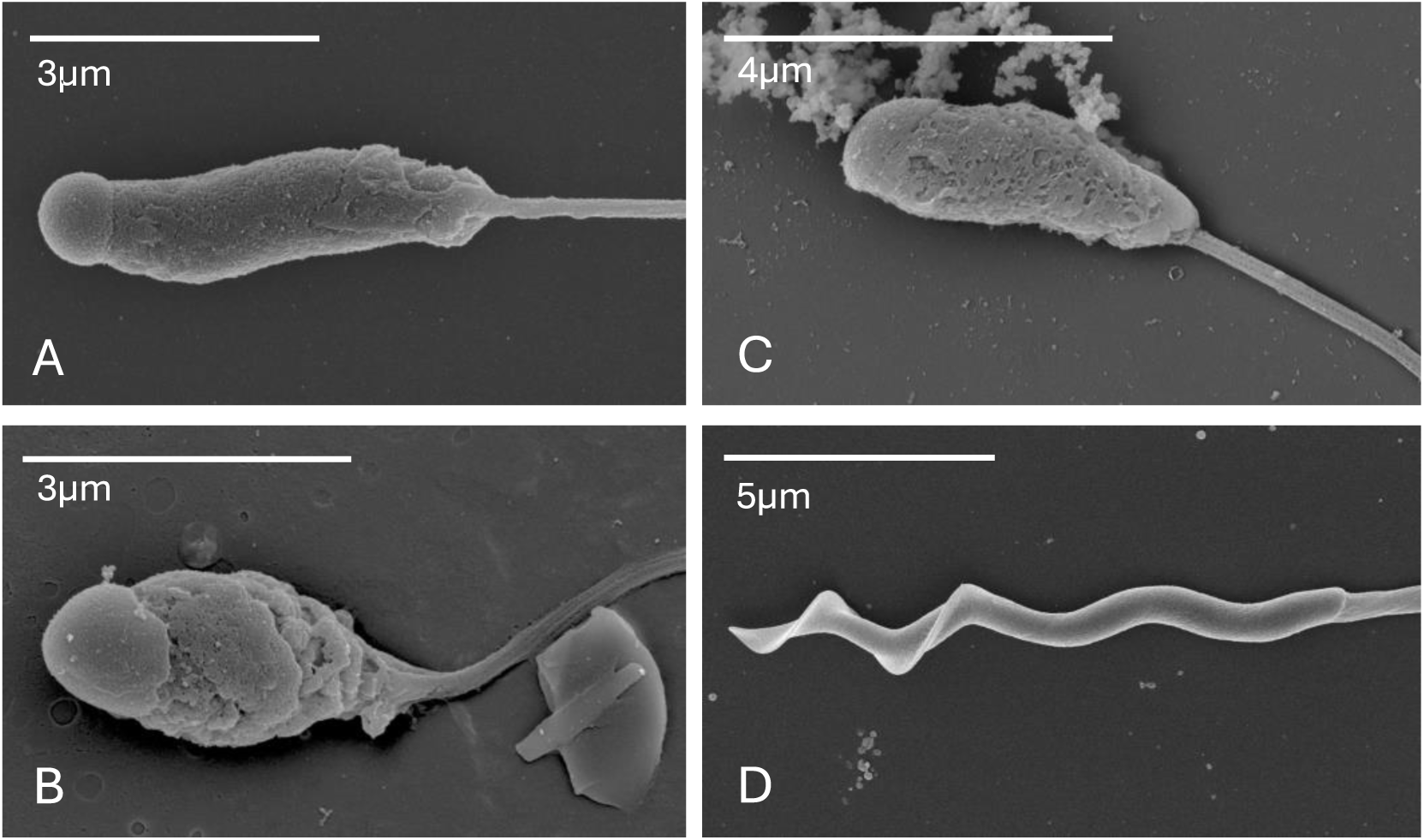
Scanning electron micrographs showing the similarity of the unusual sperm morphology in (A) the nominate subspecies of red-browed finch (*N. t. temporalis*), (B) the Eurasian bullfinch (*P. pyrrhula*), and (C) the Azores bullfinch (*P. murina*) compared to the typical passerine sperm morphology exemplified by (D) the house sparrow (*Passer domesticus*). Note that the difference in appearance of the cell surface between A and B/C is likely due to differences in fixation methods.

## Supplementary Tables

**Table S1.** Sampling locations and sampling periods.

| Species | Population | Location | Coordinates | Sampling period |
| --- | --- | --- | --- | --- |
| Red-browed finch | Lake Samsonvale | Queensland, Australia | 27°16' S, 152°51' E | November 2015 |
| Red-browed finch | Martinsville | New South Wales, Australia | 33°1'23" S, 151°23'58" E | August-December 2020 |
| Red-browed finch | Nelson | New South Wales, Australia | 33°38'39" S, 150°55'8" E | August-December 2020 |
| Red-browed finch | Narara | New South Wales, Australia | 33°24'1" S, 151°20'33" E | 2018; October 2020 |
| Red-browed finch | Burrendong Arboretum | New South Wales, Australia | 32°42'15" S, 149°6'38" E | November 2020 |
| Red-browed finch | Wellington Caves | New South Wales, Australia | 32°37'12" S, 148°56'22" E | November 2020 |
| Red-browed finch | Ballimore | New South Wales, Australia | 32°11'40" S, 148°54'0" E | November 2020 |
| Red-browed finch | NIOO-KNAW | New South Wales, Australia | 51°59'14" N, 5°40'20" E | March 2022; January 2023; April 2023 |
| Lesser red-browed finch | Tallegalla | Queensland, Australia | 27°35' 47" S, 152°34'7" E | March 2019 |
| Crimson finch | Martinsville | New South Wales, Australia | 33°1'23" S, 151°23'58" E | August 2018 |
| Double-barred finch | Wianamatta Nature Reserve | New South Wales, Australia | 33°41'39" S, 150°55'8" E | September 2017 |

**Table S2.** Whole genome sequencing (WGS) metadata.

| Study Accession | Source Voucher | IID | Species | Common Name | Sex | Material | Paired Reads | Mean Coverage | Run Accession | Publication |
| --- | --- | --- | --- | --- | --- | --- | --- | --- | --- | --- |
| PRJNA1158757 | CSIRO:B55446 | CF03 | *Neochmia phaeton* | Crimson finch | male | tissue | 135665578 | 18.6 | SRR30600842, SRR30602174 | *This study* |
| PRJNA1158757 | CSIRO:B57499 | RBF07 | *Neochmia temporalis minor* | Red-browed finch | female | tissue | 143728003 | 22.8 | SRR30600840, SRR30602172 | *This study* |
| PRJNA1158757 | CSIRO:B53194 | RBF01 | *Neochmia temporalis temporalis* | Lesser red-browed finch | male | tissue | 157220931 | 23.6 | SRR30600841, SRR30602173 | *This study* |
| PRJEB10586 | G169 | G169 | *Poephila acuticauda acuticauda* | Long-tailed finch | male | blood | 157504688 | 28.2 | ERR1013138 | Singhal et al. (2015) |
| PRJEB10586 | G183 | G183 | *Poephila acuticauda acuticauda* | Long-tailed finch | male | blood | 119402734 | 20.9 | ERR1013139 | Singhal et al. (2015) |
| PRJEB10586 | G250 | G250 | *Poephila acuticauda acuticauda* | Long-tailed finch | male | blood | 133255297 | 23.0 | ERR1013140 | Singhal et al. (2015) |
| PRJNA1158757 | CSIRO:B55429 | DBF01 | *Stizoptera bichenovii annulosa* | Double-barred finch | male | tissue | 96620540 | 14.0 | SRR30607951, SRR30610330, SRR30610331 | *This study* |
| PRJNA1158757 | *This study* | DBF11 | *Stizoptera bichenovii annulosa* | Double-barred finch | male | blood | 118327740 | 24.0 | SRR30607950 | *This study* |
| PRJEB10586 | DBF00 | DBF00 | *Stizoptera bichenovii bichenovii* | Double-barred finch | male | blood | 188213531 | 31.7 | ERR993524 | Singhal et al. (2015) |
| PRJEB10586 | 28402 | ZF02 | *Taeniopygia castanotis* | Zebra finch (Australian) | male | blood | 126359801 | 22.4 | ERR1013166 | Singhal et al. (2015) |
| PRJEB10586 | 26516 | ZF03 | *Taeniopygia castanotis* | Zebra finch (Australian) | male | blood | 161603426 | 28.7 | ERR1013167 | Singhal et al. (2015) |
| PRJEB10586 | 28404 | ZF04 | *Taeniopygia castanotis* | Zebra finch (Australian) | male | blood | 126503569 | 22.6 | ERR1013168 | Singhal et al. (2015) |
| PRJNA779480 | LSO256 | LSD02 | *Lonchura striata domestica* | Bengalese finch | male | blood | 143024354 | 31.2 | SRR16914200 | Lu et al. (2021) |
| PRJNA779480 | LDO253 | LSD04 | *Lonchura striata domestica* | Bengalese finch | male | blood | 130862907 | 28.4 | SRR16914202 | Lu et al. (2021) |
| PRJNA779480 | LDO005 | LSD05 | *Lonchura striata domestica* | Bengalese finch | male | blood | 115714789 | 25.5 | SRR16914203 | Lu et al. (2021) |

**Table S3.** Mean site-based observed heterozygosity for 30 largest chromosomes by sample.

| SPECIES.CODE | SAMPLE | CHROM | LENGTH | DP5.SITES | DP10.SITES | HET.SITES | DP5.HETEROZYGOSITY | DP10.HETEROZYGOSITY |
| --- | --- | --- | --- | --- | --- | --- | --- | --- |
| CF | CF03 | chr2 | 151653088 | 147487083 | 142150787 | 289879 | 0.001965 | 0.002039 |
| CF | CF03 | chr1 | 114020016 | 111312237 | 107215158 | 285579 | 0.002566 | 0.002664 |
| CF | CF03 | chr3 | 112598992 | 109701803 | 105721542 | 252896 | 0.002305 | 0.002392 |
| CF | CF03 | chrZ | 75396176 | 71394468 | 68428118 | 159636 | 0.002236 | 0.002333 |
| CF | CF03 | chr1A | 71569005 | 69795258 | 67076755 | 195452 | 0.002800 | 0.002914 |
| CF | CF03 | chr4 | 70982421 | 69436082 | 66879590 | 184517 | 0.002657 | 0.002759 |
| CF | CF03 | chr5 | 61663524 | 60257542 | 57792981 | 160364 | 0.002661 | 0.002775 |
| CF | CF03 | chr7 | 37979234 | 37200808 | 35687977 | 101954 | 0.002741 | 0.002857 |
| CF | CF03 | chr6 | 35592820 | 34493204 | 33032099 | 93703 | 0.002717 | 0.002837 |
| CF | CF03 | chr8 | 31251066 | 30502140 | 29164518 | 81205 | 0.002662 | 0.002784 |
| CF | CF03 | chr9 | 25529500 | 24904782 | 23799262 | 57564 | 0.002311 | 0.002419 |
| CF | CF03 | chr11 | 21108237 | 20504956 | 19513181 | 55894 | 0.002726 | 0.002864 |
| CF | CF03 | chr10 | 20543327 | 19867853 | 18928175 | 46963 | 0.002364 | 0.002481 |
| CF | CF03 | chr12 | 20384687 | 19916836 | 18995770 | 47808 | 0.002400 | 0.002517 |
| CF | CF03 | chr4A | 19491698 | 19062787 | 18182666 | 51425 | 0.002698 | 0.002828 |
| CF | CF03 | chr13 | 18052104 | 17549611 | 16636567 | 49904 | 0.002844 | 0.003000 |
| CF | CF03 | chr14 | 16340488 | 15900838 | 15074886 | 43704 | 0.002749 | 0.002899 |
| CF | CF03 | chr20 | 14749488 | 14278454 | 13459704 | 41904 | 0.002935 | 0.003113 |
| CF | CF03 | chr15 | 13771869 | 13408044 | 12638083 | 34055 | 0.002540 | 0.002695 |
| CF | CF03 | chr17 | 11257394 | 10879539 | 10171643 | 30291 | 0.002784 | 0.002978 |
| CF | CF03 | chr18 | 11152170 | 10756749 | 10029632 | 33200 | 0.003086 | 0.003310 |
| CF | CF03 | chr19 | 8335140 | 8077182 | 7593091 | 19605 | 0.002427 | 0.002582 |
| CF | CF03 | chr21 | 7924511 | 7337131 | 6707326 | 21207 | 0.002890 | 0.003162 |
| CF | CF03 | chr24 | 7620594 | 7229573 | 6619238 | 25433 | 0.003518 | 0.003842 |
| CF | CF03 | chr26 | 6804938 | 6356908 | 5733221 | 18168 | 0.002858 | 0.003169 |
| CF | CF03 | chr23 | 6804446 | 6419484 | 5830657 | 19956 | 0.003109 | 0.003423 |
| CF | CF03 | chr28 | 5866305 | 5289497 | 4756296 | 16651 | 0.003148 | 0.003501 |
| CF | CF03 | chr27 | 5378928 | 4799384 | 4235927 | 14816 | 0.003087 | 0.003498 |
| CF | CF03 | chr22 | 4608043 | 4130215 | 3627358 | 12265 | 0.002970 | 0.003381 |
| CF | CF03 | chr25 | 2880192 | 2446301 | 2031462 | 6271 | 0.002563 | 0.003087 |
| RBF | RBF07 | chr2 | 151653088 | 147713415 | 145909078 | 177997 | 0.001210 | 0.001220 |
| RBF | RBF07 | chr1 | 114020016 | 111564770 | 110236325 | 178096 | 0.001600 | 0.001620 |
| RBF | RBF07 | chr3 | 112598992 | 109956162 | 108614100 | 164811 | 0.001500 | 0.001520 |
| RBF | RBF07 | chr1A | 71569005 | 69834848 | 68892287 | 132361 | 0.001900 | 0.001920 |
| RBF | RBF07 | chr4 | 70982421 | 69511198 | 68656476 | 128386 | 0.001850 | 0.001870 |
| RBF | RBF07 | chr5 | 61663524 | 60378597 | 59510527 | 99281 | 0.001640 | 0.001670 |
| RBF | RBF07 | chr7 | 37979234 | 37261248 | 36760378 | 70744 | 0.001900 | 0.001920 |
| RBF | RBF07 | chr6 | 35592820 | 34504263 | 33998886 | 69198 | 0.002010 | 0.002040 |
| RBF | RBF07 | chr8 | 31251066 | 30542832 | 30075182 | 56722 | 0.001860 | 0.001890 |
| RBF | RBF07 | chr9 | 25529500 | 24691501 | 24310102 | 46353 | 0.001880 | 0.001910 |
| RBF | RBF07 | chr11 | 21108237 | 20566843 | 20196291 | 41144 | 0.002000 | 0.002040 |
| RBF | RBF07 | chr10 | 20543327 | 19919571 | 19581945 | 33069 | 0.001660 | 0.001690 |
| RBF | RBF07 | chr12 | 20384687 | 19955959 | 19648369 | 37219 | 0.001870 | 0.001890 |
| RBF | RBF07 | chr4A | 19491698 | 19092236 | 18802533 | 44653 | 0.002340 | 0.002370 |
| RBF | RBF07 | chr13 | 18052104 | 17585462 | 17244834 | 34732 | 0.001980 | 0.002010 |
| RBF | RBF07 | chr14 | 16340488 | 15960275 | 15664230 | 29828 | 0.001870 | 0.001900 |
| RBF | RBF07 | chr20 | 14749488 | 14318044 | 14025397 | 34139 | 0.002380 | 0.002430 |
| RBF | RBF07 | chr15 | 13771869 | 13464835 | 13190744 | 26253 | 0.001950 | 0.001990 |
| RBF | RBF07 | chr17 | 11257394 | 10934317 | 10676463 | 22246 | 0.002030 | 0.002080 |
| RBF | RBF07 | chr18 | 11152170 | 10799399 | 10519087 | 19846 | 0.001840 | 0.001890 |
| RBF | RBF07 | chr19 | 8335140 | 8100755 | 7915833 | 19337 | 0.002390 | 0.002440 |
| RBF | RBF07 | chr21 | 7924511 | 7379085 | 7080796 | 11775 | 0.001600 | 0.001660 |
| RBF | RBF07 | chr24 | 7620594 | 7252016 | 7000963 | 22584 | 0.003110 | 0.003230 |
| RBF | RBF07 | chr26 | 6804938 | 6412962 | 6121627 | 13092 | 0.002040 | 0.002140 |
| RBF | RBF07 | chr23 | 6804446 | 6480125 | 6214331 | 18082 | 0.002790 | 0.002910 |
| RBF | RBF07 | chr28 | 5866305 | 5299414 | 5004802 | 11749 | 0.002220 | 0.002350 |
| RBF | RBF07 | chr27 | 5378928 | 4883764 | 4564889 | 12396 | 0.002540 | 0.002720 |
| RBF | RBF07 | chr22 | 4608043 | 4185345 | 3904016 | 11258 | 0.002690 | 0.002880 |
| RBF | RBF07 | chr25 | 2880192 | 2506531 | 2250252 | 5352 | 0.002140 | 0.002380 |
| RBF | RBF01 | chr2 | 151653088 | 147833969 | 146116061 | 441788 | 0.002990 | 0.003020 |
| RBF | RBF01 | chr1 | 114020016 | 111630487 | 110367351 | 398881 | 0.003570 | 0.003610 |
| RBF | RBF01 | chr3 | 112598992 | 110061940 | 108799981 | 419041 | 0.003810 | 0.003850 |
| RBF | RBF01 | chrZ | 75396176 | 71250901 | 70094360 | 99204 | 0.001390 | 0.001420 |
| RBF | RBF01 | chr1A | 71569005 | 69904257 | 68992848 | 295026 | 0.004220 | 0.004280 |
| RBF | RBF01 | chr4 | 70982421 | 69566218 | 68754088 | 297442 | 0.004280 | 0.004330 |
| RBF | RBF01 | chr5 | 61663524 | 60425535 | 59586268 | 242462 | 0.004010 | 0.004070 |
| RBF | RBF01 | chr7 | 37979234 | 37280391 | 36788337 | 153490 | 0.004120 | 0.004170 |
| RBF | RBF01 | chr6 | 35592820 | 34506238 | 34025932 | 146281 | 0.004240 | 0.004300 |
| RBF | RBF01 | chr8 | 31251066 | 30538934 | 30090116 | 120533 | 0.003950 | 0.004010 |
| RBF | RBF01 | chr9 | 25529500 | 24696470 | 24336278 | 83965 | 0.003400 | 0.003450 |
| RBF | RBF01 | chr11 | 21108237 | 20568369 | 20206192 | 84250 | 0.004100 | 0.004170 |
| RBF | RBF01 | chr10 | 20543327 | 19925208 | 19593646 | 71603 | 0.003590 | 0.003650 |
| RBF | RBF01 | chr12 | 20384687 | 19942094 | 19629043 | 78747 | 0.003950 | 0.004010 |
| RBF | RBF01 | chr4A | 19491698 | 19086450 | 18803745 | 62197 | 0.003260 | 0.003310 |
| RBF | RBF01 | chr13 | 18052104 | 17597565 | 17270688 | 73232 | 0.004160 | 0.004240 |
| RBF | RBF01 | chr14 | 16340488 | 15948008 | 15656097 | 53886 | 0.003380 | 0.003440 |
| RBF | RBF01 | chr20 | 14749488 | 14299831 | 14014355 | 59060 | 0.004130 | 0.004210 |
| RBF | RBF01 | chr15 | 13771869 | 13463573 | 13197465 | 48082 | 0.003570 | 0.003640 |
| RBF | RBF01 | chr17 | 11257394 | 10934611 | 10684224 | 41613 | 0.003810 | 0.003890 |
| RBF | RBF01 | chr18 | 11152170 | 10802861 | 10520896 | 43146 | 0.003990 | 0.004100 |
| RBF | RBF01 | chr19 | 8335140 | 8117252 | 7946966 | 33274 | 0.004100 | 0.004190 |
| RBF | RBF01 | chr21 | 7924511 | 7386368 | 7087926 | 30563 | 0.004140 | 0.004310 |
| RBF | RBF01 | chr24 | 7620594 | 7263006 | 7016087 | 29444 | 0.004050 | 0.004200 |
| RBF | RBF01 | chr26 | 6804938 | 6419531 | 6134652 | 22141 | 0.003450 | 0.003610 |
| RBF | RBF01 | chr23 | 6804446 | 6471165 | 6209272 | 10322 | 0.001600 | 0.001660 |
| RBF | RBF01 | chr28 | 5866305 | 5270579 | 4974377 | 20158 | 0.003820 | 0.004050 |
| RBF | RBF01 | chr27 | 5378928 | 4875969 | 4559279 | 11122 | 0.002280 | 0.002440 |
| RBF | RBF01 | chr22 | 4608043 | 4193300 | 3912500 | 18261 | 0.004350 | 0.004670 |
| RBF | RBF01 | chr25 | 2880192 | 2523212 | 2258419 | 8345 | 0.003310 | 0.003700 |
| LTF | G169 | chr2 | 151653088 | 148771012 | 147654516 | 724971 | 0.004870 | 0.004910 |
| LTF | G183 | chr2 | 151653088 | 148352714 | 146317109 | 724340 | 0.004880 | 0.004950 |
| LTF | G250 | chr2 | 151653088 | 148618759 | 147265013 | 727321 | 0.004890 | 0.004940 |
| LTF | G169 | chr1 | 114020016 | 112213459 | 111443140 | 649704 | 0.005790 | 0.005830 |
| LTF | G183 | chr1 | 114020016 | 112045248 | 110540883 | 646527 | 0.005770 | 0.005850 |
| LTF | G250 | chr1 | 114020016 | 112149937 | 111201275 | 660286 | 0.005890 | 0.005940 |
| LTF | G169 | chr3 | 112598992 | 110601575 | 109811354 | 613724 | 0.005550 | 0.005590 |
| LTF | G183 | chr3 | 112598992 | 110281324 | 108795824 | 603505 | 0.005470 | 0.005550 |
| LTF | G250 | chr3 | 112598992 | 110446438 | 109483277 | 605164 | 0.005480 | 0.005530 |
| LTF | G169 | chrZ | 75396176 | 71924502 | 71080353 | 168999 | 0.002350 | 0.002380 |
| LTF | G183 | chrZ | 75396176 | 71665861 | 70267616 | 162570 | 0.002270 | 0.002310 |
| LTF | G250 | chrZ | 75396176 | 71786994 | 70776734 | 167802 | 0.002340 | 0.002370 |
| LTF | G169 | chr1A | 71569005 | 70305180 | 69782501 | 440212 | 0.006260 | 0.006310 |
| LTF | G183 | chr1A | 71569005 | 70170002 | 69129370 | 438334 | 0.006250 | 0.006340 |
| LTF | G250 | chr1A | 71569005 | 70238272 | 69542243 | 440208 | 0.006270 | 0.006330 |
| LTF | G169 | chr4 | 70982421 | 69980406 | 69504390 | 438768 | 0.006270 | 0.006310 |
| LTF | G183 | chr4 | 70982421 | 69840075 | 68878104 | 432933 | 0.006200 | 0.006290 |
| LTF | G250 | chr4 | 70982421 | 69917295 | 69313631 | 434813 | 0.006220 | 0.006270 |
| LTF | G169 | chr5 | 61663524 | 60773556 | 60342998 | 336793 | 0.005540 | 0.005580 |
| LTF | G183 | chr5 | 61663524 | 60650495 | 59722268 | 333668 | 0.005500 | 0.005590 |
| LTF | G250 | chr5 | 61663524 | 60713064 | 60126479 | 334184 | 0.005500 | 0.005560 |
| LTF | G169 | chr7 | 37979234 | 37507590 | 37247412 | 209490 | 0.005590 | 0.005620 |
| LTF | G183 | chr7 | 37979234 | 37404519 | 36780062 | 204847 | 0.005480 | 0.005570 |
| LTF | G250 | chr7 | 37979234 | 37455125 | 37066154 | 209838 | 0.005600 | 0.005660 |
| LTF | G169 | chr6 | 35592820 | 34836453 | 34599538 | 235261 | 0.006750 | 0.006800 |
| LTF | G183 | chr6 | 35592820 | 34729571 | 34065883 | 234540 | 0.006750 | 0.006880 |
| LTF | G250 | chr6 | 35592820 | 34792120 | 34398821 | 233185 | 0.006700 | 0.006780 |
| LTF | G169 | chr8 | 31251066 | 30789906 | 30564742 | 168962 | 0.005490 | 0.005530 |
| LTF | G183 | chr8 | 31251066 | 30706363 | 30113051 | 165980 | 0.005410 | 0.005510 |
| LTF | G250 | chr8 | 31251066 | 30742486 | 30372014 | 167303 | 0.005440 | 0.005510 |
| LTF | G169 | chr9 | 25529500 | 25169199 | 24993466 | 158944 | 0.006320 | 0.006360 |
| LTF | G183 | chr9 | 25529500 | 25099401 | 24545678 | 157295 | 0.006270 | 0.006410 |
| LTF | G250 | chr9 | 25529500 | 25137800 | 24807007 | 156033 | 0.006210 | 0.006290 |
| LTF | G169 | chr11 | 21108237 | 20756071 | 20580813 | 137840 | 0.006640 | 0.006700 |
| LTF | G183 | chr11 | 21108237 | 20690327 | 20213455 | 135830 | 0.006560 | 0.006720 |
| LTF | G250 | chr11 | 21108237 | 20721049 | 20414583 | 136099 | 0.006570 | 0.006670 |
| LTF | G169 | chr10 | 20543327 | 20107142 | 19943357 | 125135 | 0.006220 | 0.006270 |
| LTF | G183 | chr10 | 20543327 | 20010105 | 19528207 | 122350 | 0.006110 | 0.006270 |
| LTF | G250 | chr10 | 20543327 | 20050268 | 19737379 | 124070 | 0.006190 | 0.006290 |
| LTF | G169 | chr12 | 20384687 | 20131195 | 19990975 | 133014 | 0.006610 | 0.006650 |
| LTF | G183 | chr12 | 20384687 | 20074513 | 19622139 | 130216 | 0.006490 | 0.006640 |
| LTF | G250 | chr12 | 20384687 | 20098952 | 19807447 | 131575 | 0.006550 | 0.006640 |
| LTF | G169 | chr4A | 19491698 | 19251926 | 19117704 | 132384 | 0.006880 | 0.006920 |
| LTF | G183 | chr4A | 19491698 | 19197712 | 18736968 | 128474 | 0.006690 | 0.006860 |
| LTF | G250 | chr4A | 19491698 | 19220151 | 18936937 | 130845 | 0.006810 | 0.006910 |
| LTF | G169 | chr13 | 18052104 | 17772357 | 17625311 | 128380 | 0.007220 | 0.007280 |
| LTF | G183 | chr13 | 18052104 | 17696695 | 17186571 | 125893 | 0.007110 | 0.007330 |
| LTF | G250 | chr13 | 18052104 | 17728206 | 17407579 | 128553 | 0.007250 | 0.007380 |
| LTF | G169 | chr14 | 16340488 | 16134456 | 16014648 | 107284 | 0.006650 | 0.006700 |
| LTF | G183 | chr14 | 16340488 | 16077501 | 15629278 | 105761 | 0.006580 | 0.006770 |
| LTF | G250 | chr14 | 16340488 | 16096446 | 15828636 | 107861 | 0.006700 | 0.006810 |
| LTF | G169 | chr20 | 14749488 | 14468910 | 14339509 | 104112 | 0.007200 | 0.007260 |
| LTF | G183 | chr20 | 14749488 | 14405465 | 13899967 | 101498 | 0.007050 | 0.007300 |
| LTF | G250 | chr20 | 14749488 | 14430385 | 14113476 | 102302 | 0.007090 | 0.007250 |
| LTF | G169 | chr15 | 13771869 | 13581725 | 13467887 | 84194 | 0.006200 | 0.006250 |
| LTF | G183 | chr15 | 13771869 | 13518787 | 13049885 | 83876 | 0.006200 | 0.006430 |
| LTF | G250 | chr15 | 13771869 | 13543310 | 13264835 | 84521 | 0.006240 | 0.006370 |
| LTF | G169 | chr17 | 11257394 | 11072121 | 10963606 | 78327 | 0.007070 | 0.007140 |
| LTF | G183 | chr17 | 11257394 | 11013885 | 10503361 | 75712 | 0.006870 | 0.007210 |
| LTF | G250 | chr17 | 11257394 | 11035897 | 10727587 | 76379 | 0.006920 | 0.007120 |
| LTF | G169 | chr18 | 11152170 | 10925511 | 10784678 | 64857 | 0.005940 | 0.006010 |
| LTF | G183 | chr18 | 11152170 | 10856029 | 10349125 | 62806 | 0.005790 | 0.006070 |
| LTF | G250 | chr18 | 11152170 | 10881819 | 10566161 | 63822 | 0.005870 | 0.006040 |
| LTF | G169 | chr19 | 8335140 | 8243482 | 8175530 | 57790 | 0.007010 | 0.007070 |
| LTF | G183 | chr19 | 8335140 | 8211051 | 7953742 | 56008 | 0.006820 | 0.007040 |
| LTF | G250 | chr19 | 8335140 | 8219954 | 8057247 | 56753 | 0.006900 | 0.007040 |
| LTF | G169 | chr21 | 7924511 | 7517808 | 7333877 | 49666 | 0.006610 | 0.006770 |
| LTF | G183 | chr21 | 7924511 | 7399989 | 6871054 | 47833 | 0.006460 | 0.006960 |
| LTF | G250 | chr21 | 7924511 | 7447398 | 7092204 | 48682 | 0.006540 | 0.006860 |
| LTF | G169 | chr24 | 7620594 | 7384853 | 7239012 | 57635 | 0.007800 | 0.007960 |
| LTF | G183 | chr24 | 7620594 | 7299712 | 6747151 | 54669 | 0.007490 | 0.008100 |
| LTF | G250 | chr24 | 7620594 | 7331001 | 6995280 | 56409 | 0.007690 | 0.008060 |
| LTF | G169 | chr26 | 6804938 | 6527656 | 6361225 | 44936 | 0.006880 | 0.007060 |
| LTF | G183 | chr26 | 6804938 | 6406041 | 5855914 | 42823 | 0.006680 | 0.007310 |
| LTF | G250 | chr26 | 6804938 | 6455214 | 6107727 | 43539 | 0.006740 | 0.007130 |
| LTF | G169 | chr23 | 6804446 | 6603694 | 6463214 | 52007 | 0.007880 | 0.008050 |
| LTF | G183 | chr23 | 6804446 | 6507855 | 6001409 | 47623 | 0.007320 | 0.007940 |
| LTF | G250 | chr23 | 6804446 | 6547550 | 6224810 | 49657 | 0.007580 | 0.007980 |
| LTF | G169 | chr28 | 5866305 | 5519077 | 5386919 | 38745 | 0.007020 | 0.007190 |
| LTF | G183 | chr28 | 5866305 | 5444012 | 5058917 | 36337 | 0.006670 | 0.007180 |
| LTF | G250 | chr28 | 5866305 | 5472762 | 5226613 | 37933 | 0.006930 | 0.007260 |
| LTF | G169 | chr27 | 5378928 | 5013296 | 4830739 | 34590 | 0.006900 | 0.007160 |
| LTF | G183 | chr27 | 5378928 | 4905868 | 4388771 | 34033 | 0.006940 | 0.007750 |
| LTF | G250 | chr27 | 5378928 | 4956190 | 4622840 | 34780 | 0.007020 | 0.007520 |
| LTF | G169 | chr22 | 4608043 | 4333295 | 4142928 | 30542 | 0.007050 | 0.007370 |
| LTF | G183 | chr22 | 4608043 | 4194059 | 3675672 | 28052 | 0.006690 | 0.007630 |
| LTF | G250 | chr22 | 4608043 | 4244799 | 3879012 | 29013 | 0.006830 | 0.007480 |
| LTF | G169 | chr25 | 2880192 | 2670534 | 2507590 | 19010 | 0.007120 | 0.007580 |
| LTF | G183 | chr25 | 2880192 | 2540268 | 2119757 | 17252 | 0.006790 | 0.008140 |
| LTF | G250 | chr25 | 2880192 | 2589137 | 2287704 | 17533 | 0.006770 | 0.007660 |
| DBF | DBF01 | chr2 | 151653088 | 146186257 | 125286219 | 513704 | 0.003510 | 0.004100 |
| DBF | DBF11 | chr2 | 151653088 | 146284921 | 132410767 | 554317 | 0.003790 | 0.004190 |
| DBF | DBF01 | chr1 | 114020016 | 110564377 | 94541624 | 493425 | 0.004460 | 0.005220 |
| DBF | DBF11 | chr1 | 114020016 | 110356120 | 99741183 | 509347 | 0.004620 | 0.005110 |
| DBF | DBF01 | chr3 | 112598992 | 108831782 | 93106608 | 435656 | 0.004000 | 0.004680 |
| DBF | DBF11 | chr3 | 112598992 | 109023374 | 99424451 | 441418 | 0.004050 | 0.004440 |
| DBF | DBF01 | chrZ | 75396176 | 70629571 | 60351010 | 252814 | 0.003580 | 0.004190 |
| DBF | DBF11 | chrZ | 75396176 | 70516281 | 63273944 | 226657 | 0.003210 | 0.003580 |
| DBF | DBF01 | chr1A | 71569005 | 69086774 | 58920439 | 286352 | 0.004140 | 0.004860 |
| DBF | DBF11 | chr1A | 71569005 | 69210769 | 63094339 | 307465 | 0.004440 | 0.004870 |
| DBF | DBF01 | chr4 | 70982421 | 68817855 | 58805327 | 281917 | 0.004100 | 0.004790 |
| DBF | DBF11 | chr4 | 70982421 | 68899125 | 62424668 | 292147 | 0.004240 | 0.004680 |
| DBF | DBF01 | chr5 | 61663524 | 59664592 | 50190918 | 241045 | 0.004040 | 0.004800 |
| DBF | DBF11 | chr5 | 61663524 | 59911333 | 55182888 | 252430 | 0.004210 | 0.004570 |
| DBF | DBF01 | chr7 | 37979234 | 36724610 | 30725752 | 107811 | 0.002940 | 0.003510 |
| DBF | DBF11 | chr7 | 37979234 | 36907351 | 34098906 | 113652 | 0.003080 | 0.003330 |
| DBF | DBF01 | chr6 | 35592820 | 34085583 | 28244165 | 112465 | 0.003300 | 0.003980 |
| DBF | DBF11 | chr6 | 35592820 | 34359948 | 31943275 | 121974 | 0.003550 | 0.003820 |
| DBF | DBF01 | chr8 | 31251066 | 30167253 | 24927397 | 100252 | 0.003320 | 0.004020 |
| DBF | DBF11 | chr8 | 31251066 | 30389110 | 28206348 | 132534 | 0.004360 | 0.004700 |
| DBF | DBF01 | chr9 | 25529500 | 24664847 | 20140753 | 103806 | 0.004210 | 0.005150 |
| DBF | DBF11 | chr9 | 25529500 | 24919919 | 23333188 | 108978 | 0.004370 | 0.004670 |
| DBF | DBF01 | chr11 | 21108237 | 20281492 | 16572511 | 87005 | 0.004290 | 0.005250 |
| DBF | DBF11 | chr11 | 21108237 | 20520153 | 19069395 | 87054 | 0.004240 | 0.004570 |
| DBF | DBF01 | chr10 | 20543327 | 19579425 | 15934677 | 72971 | 0.003730 | 0.004580 |
| DBF | DBF11 | chr10 | 20543327 | 19922995 | 18691333 | 77629 | 0.003900 | 0.004150 |
| DBF | DBF01 | chr12 | 20384687 | 19698018 | 15915297 | 83294 | 0.004230 | 0.005230 |
| DBF | DBF11 | chr12 | 20384687 | 19931768 | 18744823 | 87941 | 0.004410 | 0.004690 |
| DBF | DBF01 | chr4A | 19491698 | 18820730 | 15133842 | 79316 | 0.004210 | 0.005240 |
| DBF | DBF11 | chr4A | 19491698 | 19066416 | 17846521 | 86109 | 0.004520 | 0.004820 |
| DBF | DBF01 | chr13 | 18052104 | 17281433 | 13649937 | 71748 | 0.004150 | 0.005260 |
| DBF | DBF11 | chr13 | 18052104 | 17607878 | 16617142 | 77429 | 0.004400 | 0.004660 |
| DBF | DBF01 | chr14 | 16340488 | 15636242 | 12430307 | 60414 | 0.003860 | 0.004860 |
| DBF | DBF11 | chr14 | 16340488 | 15959624 | 15103858 | 65586 | 0.004110 | 0.004340 |
| DBF | DBF01 | chr20 | 14749488 | 14076338 | 10897349 | 54366 | 0.003860 | 0.004990 |
| DBF | DBF11 | chr20 | 14749488 | 14363584 | 13670150 | 60371 | 0.004200 | 0.004420 |
| DBF | DBF01 | chr15 | 13771869 | 13167285 | 10187596 | 49208 | 0.003740 | 0.004830 |
| DBF | DBF11 | chr15 | 13771869 | 13497364 | 12909349 | 55187 | 0.004090 | 0.004270 |
| DBF | DBF01 | chr17 | 11257394 | 10656328 | 7886483 | 42029 | 0.003940 | 0.005330 |
| DBF | DBF11 | chr17 | 11257394 | 10964632 | 10525515 | 47740 | 0.004350 | 0.004540 |
| DBF | DBF01 | chr18 | 11152170 | 10490292 | 7824427 | 44087 | 0.004200 | 0.005630 |
| DBF | DBF11 | chr18 | 11152170 | 10805569 | 10251471 | 49381 | 0.004570 | 0.004820 |
| DBF | DBF01 | chr19 | 8335140 | 7946815 | 6061565 | 32717 | 0.004120 | 0.005400 |
| DBF | DBF11 | chr19 | 8335140 | 8138241 | 7771520 | 35385 | 0.004350 | 0.004550 |
| DBF | DBF01 | chr21 | 7924511 | 7060528 | 5077069 | 26347 | 0.003730 | 0.005190 |
| DBF | DBF11 | chr21 | 7924511 | 7382040 | 6948799 | 29161 | 0.003950 | 0.004200 |
| DBF | DBF01 | chr24 | 7620594 | 7005243 | 4856982 | 30111 | 0.004300 | 0.006200 |
| DBF | DBF11 | chr24 | 7620594 | 7255530 | 6894016 | 36296 | 0.005000 | 0.005260 |
| DBF | DBF01 | chr26 | 6804938 | 6066413 | 4055627 | 20888 | 0.003440 | 0.005150 |
| DBF | DBF11 | chr26 | 6804938 | 6461479 | 6152947 | 27882 | 0.004320 | 0.004530 |
| DBF | DBF01 | chr23 | 6804446 | 6169466 | 4183368 | 24876 | 0.004030 | 0.005950 |
| DBF | DBF11 | chr23 | 6804446 | 6519024 | 6225066 | 31433 | 0.004820 | 0.005050 |
| DBF | DBF01 | chr28 | 5866305 | 4987751 | 3396376 | 16985 | 0.003410 | 0.005000 |
| DBF | DBF11 | chr28 | 5866305 | 5388703 | 5105335 | 21629 | 0.004010 | 0.004240 |
| DBF | DBF01 | chr27 | 5378928 | 4458896 | 2939835 | 14225 | 0.003190 | 0.004840 |
| DBF | DBF11 | chr27 | 5378928 | 4904705 | 4586357 | 17926 | 0.003650 | 0.003910 |
| DBF | DBF01 | chr22 | 4608043 | 3809687 | 2417889 | 11653 | 0.003060 | 0.004820 |
| DBF | DBF11 | chr22 | 4608043 | 4175823 | 3861777 | 14665 | 0.003510 | 0.003800 |
| DBF | DBF01 | chr25 | 2880192 | 2142171 | 1191237 | 6633 | 0.003100 | 0.005570 |
| DBF | DBF11 | chr25 | 2880192 | 2544651 | 2316535 | 9537 | 0.003750 | 0.004120 |
| DBF | DBF00 | chr2 | 151653088 | 148364754 | 146582451 | 553365 | 0.003730 | 0.003780 |
| DBF | DBF00 | chr1 | 114020016 | 111954180 | 110566455 | 515337 | 0.004600 | 0.004660 |
| DBF | DBF00 | chr3 | 112598992 | 110440820 | 109213840 | 453888 | 0.004110 | 0.004160 |
| DBF | DBF00 | chrZ | 75396176 | 71735928 | 70543541 | 184340 | 0.002570 | 0.002610 |
| DBF | DBF00 | chr1A | 71569005 | 70163949 | 69349011 | 317856 | 0.004530 | 0.004580 |
| DBF | DBF00 | chr4 | 70982421 | 69852853 | 69055074 | 305275 | 0.004370 | 0.004420 |
| DBF | DBF00 | chr5 | 61663524 | 60662751 | 60047186 | 256242 | 0.004220 | 0.004270 |
| DBF | DBF00 | chr7 | 37979234 | 37339184 | 36976065 | 115307 | 0.003090 | 0.003120 |
| DBF | DBF00 | chr6 | 35592820 | 34740119 | 34448436 | 125790 | 0.003620 | 0.003650 |
| DBF | DBF00 | chr8 | 31251066 | 30738162 | 30448885 | 135629 | 0.004410 | 0.004450 |
| DBF | DBF00 | chr9 | 25529500 | 25158870 | 24963048 | 110925 | 0.004410 | 0.004440 |
| DBF | DBF00 | chr11 | 21108237 | 20737705 | 20553583 | 88527 | 0.004270 | 0.004310 |
| DBF | DBF00 | chr10 | 20543327 | 20107039 | 19938505 | 78444 | 0.003900 | 0.003930 |
| DBF | DBF00 | chr12 | 20384687 | 20120188 | 19965974 | 88626 | 0.004400 | 0.004440 |
| DBF | DBF00 | chr4A | 19491698 | 19242322 | 19091378 | 87167 | 0.004530 | 0.004570 |
| DBF | DBF00 | chr13 | 18052104 | 17762586 | 17616213 | 79205 | 0.004460 | 0.004500 |
| DBF | DBF00 | chr14 | 16340488 | 16091275 | 15976289 | 66538 | 0.004140 | 0.004160 |
| DBF | DBF00 | chr20 | 14749488 | 14489722 | 14376257 | 60658 | 0.004190 | 0.004220 |
| DBF | DBF00 | chr15 | 13771869 | 13597313 | 13507870 | 55532 | 0.004080 | 0.004110 |
| DBF | DBF00 | chr17 | 11257394 | 11041745 | 10955526 | 46412 | 0.004200 | 0.004240 |
| DBF | DBF00 | chr18 | 11152170 | 10923414 | 10803913 | 50678 | 0.004640 | 0.004690 |
| DBF | DBF00 | chr19 | 8335140 | 8209252 | 8146919 | 34799 | 0.004240 | 0.004270 |
| DBF | DBF00 | chr21 | 7924511 | 7514292 | 7359696 | 29721 | 0.003960 | 0.004040 |
| DBF | DBF00 | chr24 | 7620594 | 7353400 | 7226482 | 35255 | 0.004790 | 0.004880 |
| DBF | DBF00 | chr26 | 6804938 | 6543763 | 6420316 | 28244 | 0.004320 | 0.004400 |
| DBF | DBF00 | chr23 | 6804446 | 6596373 | 6495875 | 30977 | 0.004700 | 0.004770 |
| DBF | DBF00 | chr28 | 5866305 | 5495805 | 5388040 | 21380 | 0.003890 | 0.003970 |
| DBF | DBF00 | chr27 | 5378928 | 5047127 | 4908548 | 18403 | 0.003650 | 0.003750 |
| DBF | DBF00 | chr22 | 4608043 | 4274190 | 4130175 | 14124 | 0.003300 | 0.003420 |
| DBF | DBF00 | chr25 | 2880192 | 2640564 | 2521452 | 9442 | 0.003580 | 0.003740 |
| ZF | ZF02 | chr2 | 151653088 | 148726136 | 139308247 | 1099860 | 0.007400 | 0.007900 |
| ZF | ZF03 | chr2 | 151653088 | 149824911 | 147621854 | 1125289 | 0.007510 | 0.007620 |
| ZF | ZF04 | chr2 | 151653088 | 148985236 | 141295613 | 1105076 | 0.007420 | 0.007820 |
| ZF | ZF02 | chr1 | 114020016 | 112158901 | 104847825 | 956102 | 0.008520 | 0.009120 |
| ZF | ZF03 | chr1 | 114020016 | 112971412 | 111300922 | 980116 | 0.008680 | 0.008810 |
| ZF | ZF04 | chr1 | 114020016 | 112387541 | 106413931 | 968384 | 0.008620 | 0.009100 |
| ZF | ZF02 | chr3 | 112598992 | 110773211 | 104244599 | 875876 | 0.007910 | 0.008400 |
| ZF | ZF03 | chr3 | 112598992 | 111524618 | 109985457 | 879573 | 0.007890 | 0.008000 |
| ZF | ZF04 | chr3 | 112598992 | 110982852 | 105662269 | 873102 | 0.007870 | 0.008260 |
| ZF | ZF02 | chrZ | 75396176 | 73229245 | 68662764 | 153500 | 0.002100 | 0.002240 |
| ZF | ZF03 | chrZ | 75396176 | 73837070 | 72775717 | 132502 | 0.001790 | 0.001820 |
| ZF | ZF04 | chrZ | 75396176 | 73426031 | 69610722 | 187233 | 0.002550 | 0.002690 |
| ZF | ZF02 | chr1A | 71569005 | 70274142 | 66063260 | 555521 | 0.007910 | 0.008410 |
| ZF | ZF03 | chr1A | 71569005 | 70765089 | 69765915 | 566726 | 0.008010 | 0.008120 |
| ZF | ZF04 | chr1A | 71569005 | 70412149 | 66958480 | 560561 | 0.007960 | 0.008370 |
| ZF | ZF02 | chr4 | 70982421 | 69864804 | 65357200 | 629929 | 0.009020 | 0.009640 |
| ZF | ZF03 | chr4 | 70982421 | 70380711 | 69357579 | 638454 | 0.009070 | 0.009210 |
| ZF | ZF04 | chr4 | 70982421 | 69999082 | 66330770 | 633659 | 0.009050 | 0.009550 |
| ZF | ZF02 | chr5 | 61663524 | 60630534 | 57367495 | 615725 | 0.010160 | 0.010730 |
| ZF | ZF03 | chr5 | 61663524 | 60995171 | 60199207 | 535765 | 0.008780 | 0.008900 |
| ZF | ZF04 | chr5 | 61663524 | 60712683 | 58030592 | 614129 | 0.010120 | 0.010580 |
| ZF | ZF02 | chr7 | 37979234 | 37431339 | 35563970 | 340269 | 0.009090 | 0.009570 |
| ZF | ZF03 | chr7 | 37979234 | 37639414 | 37194229 | 344170 | 0.009140 | 0.009250 |
| ZF | ZF04 | chr7 | 37979234 | 37490307 | 35925967 | 349135 | 0.009310 | 0.009720 |
| ZF | ZF02 | chr6 | 35592820 | 34703088 | 33078802 | 355426 | 0.010240 | 0.010740 |
| ZF | ZF03 | chr6 | 35592820 | 34895466 | 34473763 | 363478 | 0.010420 | 0.010540 |
| ZF | ZF04 | chr6 | 35592820 | 34771085 | 33368771 | 357192 | 0.010270 | 0.010700 |
| ZF | ZF02 | chr8 | 31251066 | 30791348 | 29370865 | 297501 | 0.009660 | 0.010130 |
| ZF | ZF03 | chr8 | 31251066 | 30944443 | 30576625 | 298764 | 0.009650 | 0.009770 |
| ZF | ZF04 | chr8 | 31251066 | 30807372 | 29569539 | 295765 | 0.009600 | 0.010000 |
| ZF | ZF02 | chr9 | 25529500 | 25116976 | 24088365 | 252969 | 0.010070 | 0.010500 |
| ZF | ZF03 | chr9 | 25529500 | 25239907 | 24946374 | 254656 | 0.010090 | 0.010210 |
| ZF | ZF04 | chr9 | 25529500 | 25133013 | 24193031 | 252885 | 0.010060 | 0.010450 |
| ZF | ZF02 | chr11 | 21108237 | 20740293 | 19850667 | 124213 | 0.005990 | 0.006260 |
| ZF | ZF03 | chr11 | 21108237 | 20826512 | 20569878 | 124359 | 0.005970 | 0.006050 |
| ZF | ZF04 | chr11 | 21108237 | 20749103 | 19913894 | 123317 | 0.005940 | 0.006190 |
| ZF | ZF02 | chr10 | 20543327 | 20089194 | 19290715 | 186073 | 0.009260 | 0.009650 |
| ZF | ZF03 | chr10 | 20543327 | 20182561 | 19926960 | 189417 | 0.009390 | 0.009510 |
| ZF | ZF04 | chr10 | 20543327 | 20082848 | 19315071 | 188095 | 0.009370 | 0.009740 |
| ZF | ZF02 | chr12 | 20384687 | 20082222 | 19383978 | 184386 | 0.009180 | 0.009510 |
| ZF | ZF03 | chr12 | 20384687 | 20159075 | 19955199 | 185687 | 0.009210 | 0.009310 |
| ZF | ZF04 | chr12 | 20384687 | 20078638 | 19392893 | 183026 | 0.009120 | 0.009440 |
| ZF | ZF02 | chr4A | 19491698 | 19214400 | 18448087 | 199008 | 0.010360 | 0.010790 |
| ZF | ZF03 | chr4A | 19491698 | 19283801 | 19063133 | 203016 | 0.010530 | 0.010650 |
| ZF | ZF04 | chr4A | 19491698 | 19211927 | 18502411 | 198937 | 0.010350 | 0.010750 |
| ZF | ZF02 | chr13 | 18052104 | 17680601 | 17054749 | 139372 | 0.007880 | 0.008170 |
| ZF | ZF03 | chr13 | 18052104 | 17809062 | 17586344 | 194692 | 0.010930 | 0.011070 |
| ZF | ZF04 | chr13 | 18052104 | 17677704 | 17065083 | 138192 | 0.007820 | 0.008100 |
| ZF | ZF02 | chr14 | 16340488 | 16120091 | 15571435 | 123027 | 0.007630 | 0.007900 |
| ZF | ZF03 | chr14 | 16340488 | 16173638 | 15992150 | 124492 | 0.007700 | 0.007780 |
| ZF | ZF04 | chr14 | 16340488 | 16106469 | 15577639 | 123537 | 0.007670 | 0.007930 |
| ZF | ZF02 | chr20 | 14749488 | 14497644 | 14062751 | 153339 | 0.010580 | 0.010900 |
| ZF | ZF03 | chr20 | 14749488 | 14549400 | 14385886 | 154556 | 0.010620 | 0.010740 |
| ZF | ZF04 | chr20 | 14749488 | 14486513 | 14049961 | 152726 | 0.010540 | 0.010870 |
| ZF | ZF02 | chr15 | 13771869 | 13563920 | 13161667 | 132361 | 0.009760 | 0.010060 |
| ZF | ZF03 | chr15 | 13771869 | 13604167 | 13451144 | 133969 | 0.009850 | 0.009960 |
| ZF | ZF04 | chr15 | 13771869 | 13549633 | 13140201 | 133115 | 0.009820 | 0.010130 |
| ZF | ZF02 | chr17 | 11257394 | 11053687 | 10745002 | 110011 | 0.009950 | 0.010240 |
| ZF | ZF03 | chr17 | 11257394 | 11082539 | 10948629 | 111305 | 0.010040 | 0.010170 |
| ZF | ZF04 | chr17 | 11257394 | 11048768 | 10728047 | 110030 | 0.009960 | 0.010260 |
| ZF | ZF02 | chr18 | 11152170 | 10891741 | 10500374 | 90640 | 0.008320 | 0.008630 |
| ZF | ZF03 | chr18 | 11152170 | 10939680 | 10769548 | 91533 | 0.008370 | 0.008500 |
| ZF | ZF04 | chr18 | 11152170 | 10893644 | 10483624 | 90426 | 0.008300 | 0.008630 |
| ZF | ZF02 | chr19 | 8335140 | 8198590 | 7947260 | 82165 | 0.010020 | 0.010340 |
| ZF | ZF03 | chr19 | 8335140 | 8223252 | 8124275 | 82562 | 0.010040 | 0.010160 |
| ZF | ZF04 | chr19 | 8335140 | 8194027 | 7943140 | 81935 | 0.010000 | 0.010320 |
| ZF | ZF02 | chr21 | 7924511 | 7476367 | 7100667 | 73219 | 0.009790 | 0.010310 |
| ZF | ZF03 | chr21 | 7924511 | 7547820 | 7340676 | 74068 | 0.009810 | 0.010090 |
| ZF | ZF04 | chr21 | 7924511 | 7463212 | 7065799 | 73540 | 0.009850 | 0.010410 |
| ZF | ZF02 | chr24 | 7620594 | 7441553 | 7182322 | 77769 | 0.010450 | 0.010830 |
| ZF | ZF03 | chr24 | 7620594 | 7481724 | 7353390 | 80739 | 0.010790 | 0.010980 |
| ZF | ZF04 | chr24 | 7620594 | 7438074 | 7153265 | 77972 | 0.010480 | 0.010900 |
| ZF | ZF02 | chr26 | 6804938 | 6484505 | 6188199 | 63081 | 0.009730 | 0.010190 |
| ZF | ZF03 | chr26 | 6804938 | 6603310 | 6424884 | 71841 | 0.010880 | 0.011180 |
| ZF | ZF04 | chr26 | 6804938 | 6471265 | 6149372 | 62680 | 0.009690 | 0.010190 |
| ZF | ZF02 | chr23 | 6804446 | 6590489 | 6322801 | 51323 | 0.007790 | 0.008120 |
| ZF | ZF03 | chr23 | 6804446 | 6636841 | 6494942 | 52824 | 0.007960 | 0.008130 |
| ZF | ZF04 | chr23 | 6804446 | 6587393 | 6288464 | 51748 | 0.007860 | 0.008230 |
| ZF | ZF02 | chr28 | 5866305 | 5461535 | 5158343 | 54757 | 0.010030 | 0.010620 |
| ZF | ZF03 | chr28 | 5866305 | 5523496 | 5352529 | 54689 | 0.009900 | 0.010220 |
| ZF | ZF04 | chr28 | 5866305 | 5452059 | 5134554 | 54962 | 0.010080 | 0.010700 |
| ZF | ZF02 | chr27 | 5378928 | 4985303 | 4611023 | 47209 | 0.009470 | 0.010240 |
| ZF | ZF03 | chr27 | 5378928 | 5015234 | 4787253 | 46066 | 0.009190 | 0.009620 |
| ZF | ZF04 | chr27 | 5378928 | 4934488 | 4530264 | 46560 | 0.009440 | 0.010280 |
| ZF | ZF02 | chr22 | 4608043 | 4331626 | 4022285 | 37869 | 0.008740 | 0.009410 |
| ZF | ZF03 | chr22 | 4608043 | 4402460 | 4222530 | 40025 | 0.009090 | 0.009480 |
| ZF | ZF04 | chr22 | 4608043 | 4319717 | 3990425 | 38456 | 0.008900 | 0.009640 |
| ZF | ZF02 | chr25 | 2880192 | 2583029 | 2305108 | 25942 | 0.010040 | 0.011250 |
| ZF | ZF03 | chr25 | 2880192 | 2649630 | 2462572 | 27360 | 0.010330 | 0.011110 |
| ZF | ZF04 | chr25 | 2880192 | 2568753 | 2252885 | 25931 | 0.010090 | 0.011510 |

**Table S4.** Summary of mean site-based observed heterozygosity by species.

| Species | Common Name | N | Autosomal Heterozygosity | ChrZ Heterozygosity | Z:A Heterozygosity |
| --- | --- | --- | --- | --- | --- |
| *Taeniopygia guttata castanotis* | Zebra Finch | 3 | 0.00902 | 0.00225 | 0.250 |
| *Poephila acuticauda acuticauda* | Long-tailed Finch | 3 | 0.00600 | 0.00235 | 0.392 |
| *Stizoptera bichenovii bichenovii* | Double-barred Finch | 1 | 0.00420 | 0.00261 | 0.621 |
| *Stizoptera bichenovii annulosa* | Double-barred Finch | 2 | 0.00460 | 0.00389 | 0.845 |
| *Neochmia phaeton* | Crimson Finch | 1 | 0.00264 | 0.00233 | 0.883 |
| *Neochmia temporalis temporalis* | Red-browed Finch | 1 | 0.00379 | 0.00142 | 0.375 |
| *Neochmia temporalis minor* | Red-browed Finch | 1 | 0.00175 | N/A | N/A |

Mean site-based autosomal observed heterozygosity calculated as the number of heterozygous sites / number of sites with DP >=10.
Heterozygosity calculated from 29 largest chromosomes >2.7 Mb and reported as mean for each species
Data only from a single female for *Neochmia temporalis minor*

## Literature cited

Aire, T. A. 2014. Spermiogenesis in birds. Spermatogenesis 4:e959392.

Aoki, V. W., L. Liu, and D. T. Carrell. 2005. Identification and evaluation of a novel sperm protamine abnormality in a population of infertile males. Human Reproduction 20:1298–1306.

Arévalo, L., G. E. Merges, S. Schneider, F. E. Oben, I. S. Neumann, and H. Schorle. 2022. Loss of the cleaved-protamine 2 domain leads to incomplete histone-to-protamine exchange and infertility in mice. PLOS Genetics 18:e1010272. Public Library of Science.

Ausió, J., J. T. Soley, W. Burger, J. D. Lewis, D. Barreda, and K. M. Cheng. 1999. The Histidine-Rich Protamine from Ostrich and Tinamou Sperm. A Link between Reptile and Bird Protamines. Biochemistry 38:180–184.

Bartolomaeus, T., V. Bronkars, L. Adam, and J. von Döhren. 2024. Ultrastructure and phylogenetic significance of spermatozoa in Nemertea. Zoomorphology, doi: 10.1007/s00435-024-00659-2.

Bergeron, L. A., S. Besenbacher, J. Zheng, P. Li, M. F. Bertelsen, B. Quintard, J. I. Hoffman, Z. Li, J. St. Leger, C. Shao, J. Stiller, M. T. P. Gilbert, M. H. Schierup, and G. Zhang. 2023. Evolution of the germline mutation rate across vertebrates. Nature 615:285–291. Nature Publishing Group.

Bird, J. P., R. Martin, H. R. Akçakaya, J. Gilroy, I. J. Burfield, S. T. Garnett, A. Symes, J. Taylor, Ç. H. Şekercioğlu, and S. H. M. Butchart. 2020. Generation lengths of the world’s birds and their implications for extinction risk. Conservation Biology 34:1252–1261.

Birkhead, T. R., F. Giusti, S. Immler, and B. G. M. Jamieson. 2007. Ultrastructure of the unusual spermatozoon of the Eurasion bullfinch (Pyrrhula pyrrhula). Acta Zool 88:119–128.

Birkhead, T. R., and S. Immler. 2006. Making sperm: design, quality control and sperm competition. Society of Reproduction and Fertility Supplement 1–9.

Birkhead, T. R., S. Immler, E. J. Pellatt, and R. Freckleton. 2006. Unusual sperm morphology in the eurasian bullfinch (Pyrrhula pyrrhula). The Auk 123:383–392. University of California Press on behalf of the American Ornithologists’ Union.

Birkhead, T. R., and R. Montgomerie. 2009. Three centuries of sperm research. Pp. 1–42 in Sperm Biology: An Evolutionary Perspective. Academic Press, San Diego.

Birkhead, T. R., E. J. Pellatt, P. Brekke, R. Yeates, and H. Castillo-Juarez. 2005. Genetic effects on sperm design in the zebra finch. Nature 434:383–387. Nature Publishing Group.

Breed, W. G. 1993. Novel organization of the spermatozoon in two species of murid rodents from Southern Asia. Reproduction 99:149–158. Bioscientifica Ltd.

Breed, W. G. 1995. Spermatozoa of murid rodents from Africa: morphological diversity and evolutionary trends. Journal of Zoology 237:625–651.

Breed, W. G. 1997. Unusual chromatin structural organization in the sperm head of a murid rodent from Southern Africa: the red veld rat, Aethomys chrysophilus type B. Reproduction 111:221–228. Bioscientifica Ltd.

Breed, W. G. 1983. Variation in sperm morphology in the Australian rodent genus, Pseudomys (Muridae). Cell Tissue Res. 229:611–625.

Breed, W. G., M. Bauer, R. Wade, N. Thitipramote, J. Suwajarat, and L. Yelland. 2007. Intra-individual variation in sperm tail length in murine rodents. Journal of Zoology 272:299–304.

Brouwer, L., and S. C. Griffith. 2019. Extra-pair paternity in birds. Molecular Ecology 9:855–19. Blackwell Science Ltd.

Callaghan, C. T., S. Nakagawa, and W. K. Cornwell. 2021. Global abundance estimates for 9,700 bird species. PNAS 118. National Academy of Sciences.

Carrell, D. T., and L. Liu. 2001. Altered Protamine 2 Expression Is Uncommon In Donors of Known Fertility, but Common Among Men With Poor Fertilizing Capacity, and May Reflect Other Abnormalities of Spermiogenesis. Journal of Andrology 22:604–610.

Chiva, M., H. F. Kasinsky, and J. A. Subirana. 1987. Characterization of protamines from four avian species. FEBS Letters 215:237–240.

Cho, C., H. Jung-Ha, W. D. Willis, E. H. Goulding, P. Stein, Z. Xu, R. M. Schultz, N. B. Hecht, and E. M. Eddy. 2003. Protamine 2 Deficiency Leads to Sperm DNA Damage and Embryo Death in Mice1. Biology of Reproduction 69:211–217.

Christidis, L. 1987. Chromosomal evolution within the family Estrildidae (Aves) III. The Estrildae (waxbill finches). Genetica 72:93–100.

Danecek, P., A. Auton, G. Abecasis, C. A. Albers, E. Banks, M. A. DePristo, R. E. Handsaker, G. Lunter, G. T. Marth, S. T. Sherry, G. McVean, R. Durbin, and 1000 Genomes Project Analysis Group. 2011. The variant call format and VCFtools. Bioinformatics 27:2156–2158.

Danecek, P., J. K. Bonfield, J. Liddle, J. Marshall, V. Ohan, M. O. Pollard, A. Whitwham, T. Keane, S. A. McCarthy, R. M. Davies, and H. Li. 2021. Twelve years of SAMtools and BCFtools. GigaScience 10:giab008.

Durrant, K. L., D. A. Dawson, T. Burke, and T. R. Birkhead. 2010. The Unusual Sperm Morphology of the Eurasian Bullfinch (Pyrrhula pyrrhula) is not Due to the Phenotypic Result of Genetic Reduction. The Auk 127:832–840.

Fitzpatrick, J. L., A. F. Kahrl, and R. R. Snook. 2022. SpermTree, a species-level database of sperm morphology spanning the animal tree of life. Sci Data 9:30. Nature Publishing Group.

Flynn, J. M., R. Hubley, C. Goubert, J. Rosen, A. G. Clark, C. Feschotte, and A. F. Smit. 2020. RepeatModeler2 for automated genomic discovery of transposable element families. Proceedings of the National Academy of Sciences 117:9451–9457. National Acad Sciences.

Franzén, Å. 1956. On spermiogenesis, morphology of the spermatozoon, and biology of fertilization among invertebrates. Zool. Bidrag. Uppsala 31:355–482.

Gage, M. J. G., A. K. Surridge, J. L. Tomkins, E. Green, L. Wiskin, D. J. Bell, and G. M. Hewitt. 2006. Reduced Heterozygosity Depresses Sperm Quality in Wild Rabbits, Oryctolagus cuniculus. Current Biology 16:612–617. Elsevier.

Good, B. H., and M. M. Desai. 2014. Deleterious Passengers in Adapting Populations. Genetics 198:1183–1208.

Hilgers, L., S. Liu, A. Jensen, T. Brown, T. Cousins, R. Schweiger, K. Guschanski, and M. Hiller. 2024. Avoidable false PSMC population size peaks occur across numerous studies. bioRxiv.

Hooper, D. M., and T. D. Price. 2017. Chromosomal inversion differences correlate with range overlap in passerine birds. Nat Ecol Evol 1:1526–1534. Nature Publishing Group.

Hooper, D. M., and T. D. Price. 2015. Rates of karyotypic evolution in Estrildid finches differ between island and continental clades. Evolution 69:890–903. Society for the Study of Evolution.

Immler, S., S. C. Griffth, R. Zann, and T. R. Birkhead. 2012. Intra-specific variance in sperm morphometry: a comparison between wild and domesticated Zebra Finches Taeniopygia guttata. Ibis 154:480–487.

Jamieson, B. G. M. 2007. Avian spermatozoa: structure and phylogeny. Pp. 349–511 in B. G. M. Jamieson, ed. Reproductive biology and phylogeny of birds. Enfield, NH.

Jamieson, B. G. M. 1987. The ultrastructure and phylogeny of insect spermatozoa. Cambridge University Press.

Jamieson, B. G. M., J. Ausio, and J.-L. Justine. 1995. Advances in Spermatozoal Phylogeny and Taxonomy. Muséum national d’Histoire naturelle, Paris.

Kahrl, A. F., R. R. Snook, and J. L. Fitzpatrick. 2022. Fertilization mode differentially impacts the evolution of vertebrate sperm components. Nat Commun 13:6809. Nature Publishing Group.

Kahrl, A. F., R. R. Snook, and J. L. Fitzpatrick. 2021. Fertilization mode drives sperm length evolution across the animal tree of life. Nat Ecol Evol, doi: 10.1038/s41559-021-01488-y.

Kim, K.-W., C. Bennison, N. Hemmings, L. Brookes, L. L. Hurley, S. C. Griffith, T. Burke, T. R. Birkhead, and J. Slate. 2017. A sex-linked supergene controls sperm morphology and swimming speed in a songbird. Nat. ecol. evol. 1–9. Springer US.

Knief, U., W. Forstmeier, Y. Pei, M. Ihle, D. Wang, K. Martin, P. Opatová, J. Albrechtová, M. Wittig, A. Franke, T. Albrecht, and B. Kempenaers. 2017. A sex-chromosome inversion causes strong overdominance for sperm traits that affect siring success. Nat. ecol. evol. 1:1177–1184. Springer US.

Kucera, A. C., and B. J. Heidinger. 2018. Avian Semen Collection by Cloacal Massage and Isolation of DNA from Sperm. JoVE (Journal of Visualized Experiments) e55324.

Laskemoen, T., O. Kleven, F. Fossøy, and J. T. Lifjeld. 2007. Intraspecific variation in sperm length in two passerine species, the Bluethroat Luscinia svecica and the Willow Warbler Phylloscopus trochilus. 84:9.

Li, H. 2013. Aligning sequence reads, clone sequences and assembly contigs with BWA-MEM. arXiv.

Li, H., and R. Durbin. 2011. Inference of human population history from individual whole-genome sequences. Nature 475:493–496. Nature Publishing Group.

Lifjeld, J. T. 2019. The avian sperm collection in the Natural History Museum, University of Oslo. Alauda 87:93–101.

Lifjeld, J. T., A. Hoenen, L. E. Johannessen, T. Laskemoen, R. J. Lopes, P. Rodrigues, and M. Rowe. 2013. The Azores bullfinch (Pyrrhula murina) has the same unusual and size-variable sperm morphology as the Eurasian bullfinch (Pyrrhula pyrrhula). Biological Journal of the Linnean Society 108:677–687. Blackwell Publishing Ltd.

Lu, C.-W., C.-T. Yao, and C.-M. Hung. 2022. Domestication obscures genomic estimates of population history. Molecular Ecology 31:752–766.

Lüke, L., A. Vicens, M. Tourmente, and E. R. S. Roldan. 2014. Evolution of Protamine Genes and Changes in Sperm Head Phenotype in Rodents. Biology of Reproduction, doi: 10.1095/biolreprod.113.115956.

Lüpold, S., R. A. de Boer, J. P. Evans, J. L. Tomkins, and J. L. Fitzpatrick. 2020. How sperm competition shapes the evolution of testes and sperm: a meta-analysis. Philosophical Transactions of the Royal Society B: Biological Sciences 375:20200064. Royal Society.

Lupold, S., and S. Pitnick. 2018. Sperm form and function: what do we know about the role of sexual selection? REPRODUCTION 155:R229–R243.

Martin, M. 2015. Cutadapt removes adapter sequences from high-throughput sequencing reads. EMBnet.journal 17:1–3.

Maynard-Smith, J., and J. Haigh. 1974. The hitch-hiking effect of a favourable gene. Genet. Res. 23:23–35.

McCarthy, E., C. S. McDiarmid, L. L. Hurley, M. Rowe, and S. C. Griffith. 2021. Highly variable sperm morphology in the masked finch (Poephila personata) and other estrildid finches. Biological Journal of the Linnean Society, doi: 10.1093/biolinnean/blab048.

McDiarmid, C. S., R. Li, A. F. Kahrl, M. Rowe, and S. C. Griffith. 2021. Sperm Sizer: a program to semi-automate the measurement of sperm length. Behav Ecol Sociobiol 75:84.

Møller, A. P. 1991. Sperm competition, sperm depletion, paternal care, and relative testis size in birds. Am Nat 137:882–906. The University of Chicago Press.

Møller, A. P., and J. V. Briskie. 1995. Extra-pair paternity, sperm competition and the evolution of testis size in birds. Behav Ecol Sociobiol 36:357–365.

Morcombe, M. K. 2003. Field guide to Australian birds. Steve Parishh Publishing, Archerfield, Qld.

Nakano, M., T. Tobita, and T. Ando. 1976. Studies on a Protamine (galline) from Fowl Sperm. International Journal of Peptide and Protein Research 8:579–587.

Oliva, R., and G. H. Dixon. 1989. Chicken Protamine Genes Are Intronless. Journal of Biological Chemistry 264:12472–12481.

Olsson, U., and P. Alström. 2020. A comprehensive phylogeny and taxonomic evaluation of the waxbills (Aves: Estrildidae). Molecular Phylogenetics and Evolution 146:106757.

Omotoriogun, T. C., T. Albrecht, J. Gohli, D. Hořák, L. E. Johannessen, A. Johnsen, J. Kreisinger, P. Z. Marki, U. Ottosson, M. Rowe, O. Sedláček, and J. T. Lifjeld. 2020. Sperm length variation among Afrotropical songbirds reflects phylogeny rather than adaptations to the tropical environment. Zoology 140:125770. Elsevier.

Payne, R. B. 2020. Red-browed Firetail (Neochmia temporalis). P. in Birds of the World (J. del Hoyo, A. Elliott, J. Sargatal, D. A. Christie, and E. de Juana, Editors). Cornell Lab of Ornithology, Ithaca, NY, USA.

Pitnick, S., D. J. Hosken, and T. R. Birkhead. 2009. Sperm Morphological Diversity. Pp. 247–394 in T. R. Birkhead, D. J. Hosken, and S. Pitnick, eds. Sperm Biology. London.

Posit team. 2023. RStudio: Integrated Development for R. Posit Software, PBC, Boston, MA.

R Core Team. 2024. R: A Language and Environment for Statistical Computing}. R Foundation for Statistical Computing, Vienna, Austria.

Rhie, A., S. A. McCarthy, O. Fedrigo, J. Damas, G. Formenti, S. Koren, M. Uliano-Silva, W. Chow, A. Fungtammasan, J. Kim, and others. 2021. Towards complete and error-free genome assemblies of all vertebrate species. Nature 592:737–746. Nature Publishing Group.

Roldan, E., and M. Gomendio. 2009. Sperm and conservation. Pp. 539–564 in Sperm Biology. Academic Press, London.

Rowe, M., T. Albrecht, E. R. A. Cramer, A. Johnsen, T. Laskemoen, J. T. Weir, and J. T. Lifjeld. 2015a. Postcopulatory sexual selection is associated with accelerated evolution of sperm morphology. Evolution 69:1044–1052.

Rowe, M., S. C. Griffith, A. Hofgaard, and J. T. Lifjeld. 2015b. Subspecific variation in sperm morphology and performance in the Long-tailed Finch (Poephila acuticauda). Avian Res 6:23. BioMed Central.

Rule, S., B. W. Brook, S. G. Haberle, C. S. M. Turney, A. P. Kershaw, and C. N. Johnson. 2012. The Aftermath of Megafaunal Extinction: Ecosystem Transformation in Pleistocene Australia. Science 335:1483–1486. American Association for the Advancement of Science.

Segami, J. C., M. Semon, C. Cunha, C. Bergin, C. F. Mugal, and A. Qvarnström. 2022. Single-Cell Transcriptomics reveals relaxed evolutionary constraint of spermatogenesis in two passerine birds as compared to mammals. bioRxiv.

Simmons, L. W., and J. L. Fitzpatrick. 2012. Sperm wars and the evolution of male fertility. Reproduction 144:519–534.

Singhal, S., E. Leffler, K. Sannareddy, I. Turner, O. Venn, D. Hooper, A. Strand, Q. Li, B. Raney, C. Balakrishnan, S. Griffith, G. McVean, and M. Przeworski. 2015. Stable recombination hotspots in birds. 1–16.

Snook, R. R. 2005. Sperm in competition: not playing by the numbers. Trends in Ecology & Evolution 20:46–53.

Sokal, R. R., and F. J. Rohlf. 1981. Biometry. W.H. Freeman and Company, San Francisco, California.

Stamatakis, A. 2014. RAxML version 8: a tool for phylogenetic analysis and post-analysis of large phylogenies. Bioinformatics 30:1312–1313.

Støstad, H. N., A. Johnsen, J. T. Lifjeld, and M. Rowe. 2018. Sperm head morphology is associated with sperm swimming speed: A comparative study of songbirds using electron microscopy. Evolution 72:1918–1932.

Supriya, K., M. Rowe, T. Laskemoen, D. Mohan, T. D. Price, and J. T. Lifjeld. 2016. Early diversification of sperm size in the evolutionary history of the old world leaf warblers (Phylloscopidae). Journal of Evolutionary Biology 29:777–789.

Thitipramote, N., J. Suwanjarat, C. Leigh, and W. G. Breed. 2011. Variation in sperm morphology of a murine rodent from South-East Asia: the Greater Bandicoot Rat, Bandicota indica. Acta Zoologica 92:201–205.

Töpfer, T., E. Haring, T. R. Birkhead, R. J. Lopes, L. L. Severinghaus, J. Martens, and M. Päckert. 2011. A molecular phylogeny of bullfinches Pyrrhula Brisson, 1760 (Aves: Fringillidae). Molecular Phylogenetics and Evolution 58:271–282.

van der Auwera, G., and B. D. O’Connor. 2020. Genomics in the Cloud: Using Docker, GATK, and WDL in Terra. O’Reilly Media, Incorporated.

van der Horst, G., L. Maree, S. H. Kotzé, and M. J. O’Riain. 2011. Sperm structure and motility in the eusocial naked mole-rat, Heterocephalus glaber: a case of degenerative orthogenesis in the absence of sperm competition? BMC Evolutionary Biology 11:351.

## Literature cited

Cramer, E. R. A., M. Rowe, F. Eroukhmanoff, J. T. Lifjeld, G.-P. Sætre, and A. Johnsen. 2019. Measuring sperm swimming performance in birds: effects of dilution, suspension medium, mechanical agitation, and sperm number. J Ornithol 160:1053–1063.

Kleven, O., F. Fossøy, T. Laskemoen, R. J. Robertson, G. Rudolfsen, and J. T. Lifjeld. 2009. Comparative Evidence for the Evolution of Sperm Swimming Speed by Sperm Competition and Female Sperm Storage Duration in Passerine Birds. Evolution 63:2466–2473.

